# Cannabis smoke and oral Δ9THC enhance working memory in aged but not young adult subjects

**DOI:** 10.1101/2024.09.26.615028

**Authors:** Sabrina Zequeira, Emely A Gazarov, Alara A Guvenli, Erin C Berthold, Alexandria S Senetra, Marcelo Febo, Takato Hiranita, Lance R McMahon, Abhisheak Sharma, Christopher R McCurdy, Barry Setlow, Jennifer L Bizon

## Abstract

With increased legalization of recreational and medical cannabis, use of this drug is growing rapidly among older adults. As cannabis use can impair cognition in young adults, it is critically important to understand how consumption interacts with the cognitive profile of aged individuals, who are already at increased risk of decline. The current study was designed to determine how cannabis influences multiple forms of cognition in young adult and aged rats of both sexes when delivered via two translationally-relevant routes of administration. Acute exposure to cannabis smoke enhanced prefrontal cortex-dependent working memory accuracy in aged males, but impaired accuracy in aged females, while having no effects in young adults of either sex. In contrast, the same cannabis smoke exposure regimen had minimal effects on a hippocampus-dependent trial-unique non-matching to location mnemonic task, irrespective of age or sex. In a second set of experiments, chronic oral consumption of Δ9-tetrahydrocannabinol (Δ9THC) enhanced working memory in aged rats of both sexes, while having no effects in young adults. In contrast, the same oral Δ9THC regimen did not affect spatial learning and memory in either age group. Minimal age differences were observed in Δ9THC pharmacokinetics with either route of administration. Together, these results show that cannabis and Δ9THC can attenuate working memory impairments that emerge in aging. While these enhancing effects do not extend to hippocampus-dependent cognition, cannabis does not appear to exacerbate age-associated impairments in this cognitive domain.

## INTRODUCTION

Cannabis use among older adults is growing rapidly, with estimates from the National Survey on Drug Use and Health showing that from 2015 to 2023 the prevalence of past-year use among adults over age 65 rose from 2.4 to 6.9% [1–4]. Many of these individuals report regular use, and among adults ages 50-64 who use cannabis, 34% of women and 39% of men report daily or near-daily use [5,6]. Despite the growing number of older cannabis users, most studies investigating effects of cannabis on cognition have been restricted to young adult subjects.

Older adults exhibit impairments in multiple forms of cognition, including executive functions such as working memory, as well as spatial and episodic/declarative memory [7–12]. These age-associated cognitive impairments can be modeled in rats. Compared to young adults, aged rats show declines in working memory tasks dependent upon prefrontal cortex (PFC) [13–17], as well as in spatial and episodic memory tasks dependent upon the hippocampus and medial temporal lobe [18–22]. Given that cannabis and cannabinoids can impair these same aspects of cognition [23–27], as well as the fact that other drugs with abuse liability (e.g., alcohol and opioids) have greater adverse effects in older individuals [28–31], there is concern that cannabis use in older adults could exacerbate existing cognitive vulnerabilities. In contrast to these concerns, however, a few studies suggest that cannabis may have fewer adverse cognitive effects and even provide some benefit in older compared to younger adults [32,33]. In addition, a few studies show that chronic administration of the primary psychoactive component of cannabis, Δ9-tetrahydrocannabinol (Δ9THC), via osmotic mini-pump enhanced performance of aged male mice across several cognitive tasks, while having no effects or even impairing performance in young adults [34,35]. These results suggest possible benefits of Δ9THC in aged subjects; however, the passive drug delivery in this prior work did not mirror either the temporal dynamics or routes of administration of human cannabis use, which can influence its cognitive consequences via differences in the speed of absorption and/or metabolism [36].

The current study was designed to more fully elucidate the cognitive effects of cannabis delivered through multiple routes of administration and time courses that mimic those frequently used by older adults (acute intake of cannabis smoke and repeated voluntary oral consumption of Δ9THC). Moreover, the study included both male and female subjects (to our knowledge, the first preclinical aging study to date examining sex-specific effects of cannabis use on cognition). The cognitive assessments were designed to independently evaluate effects of cannabis and Δ9THC on PFC - and hippocampus-dependent cognition. To assess PFC-dependent working memory, a delayed response task dependent on the medial PFC but not hippocampus was used [37]. In contrast, acute effects of cannabis smoke were assessed on a trial unique non-match to location (TUNL) task sensitive to damage to the hippocampus but not PFC [38,39]. For the repeated-exposure oral Δ9THC consumption study, the water maze was used as a “gold standard” assessment of hippocampal-dependent cognition. Parallel pharmacokinetic studies were performed to aid in the interpretation of age- and sex-dependent effects of cannabis on cognition. The results show that both acute cannabis smoke inhalation and repeated oral Δ9THC consumption can enhance cognition in aged rats, but that these enhancing effects are sex- and cognitive domain-specific.

## RESULTS

### Experiment 1: Effects of acute cannabis smoke on cognition in young adult and aged rats

In separate experiments, young adult and aged rats of both sexes were trained in touchscreen operant chambers on either a delayed response working memory task (Experiment 1a) or the TUNL task (Experiment 1b). Upon reaching stable baseline performance, rats were tested in the tasks immediately following acute exposure to smoke from 0, 3, and 5 consecutively-burned cannabis cigarettes (NIDA Drug Supply Program, ∼700 mg, 5.6% Δ9THC) as in [40–42], using a randomized, within-subjects design such that each rat was tested in separate sessions under each exposure condition, with at least 48 h between exposure sessions.

#### Experiment 1a. Delayed non-match to sample working memory task

The delayed response working memory task [13,37] was composed of multiple, self-paced trials, each consisting of a sample, delay, and choice phase. During the sample phase, one panel on the touchscreen (far left or far right, randomized across trials) was illuminated (Figure 1A). Touching the illuminated panel caused the screen to go blank and initiated the delay phase (0, 6, 12, or 24 s), during which the food trough was illuminated and the rat had to nosepoke into the trough. After the delay, both the sample and the panel on the opposite side of the screen were illuminated. A touch on the panel that was not illuminated during the sample phase (i.e., a “non-match”) yielded a food pellet. A touch on the “incorrect” panel caused the screen to go blank and no food was delivered.

**Figure 1.**
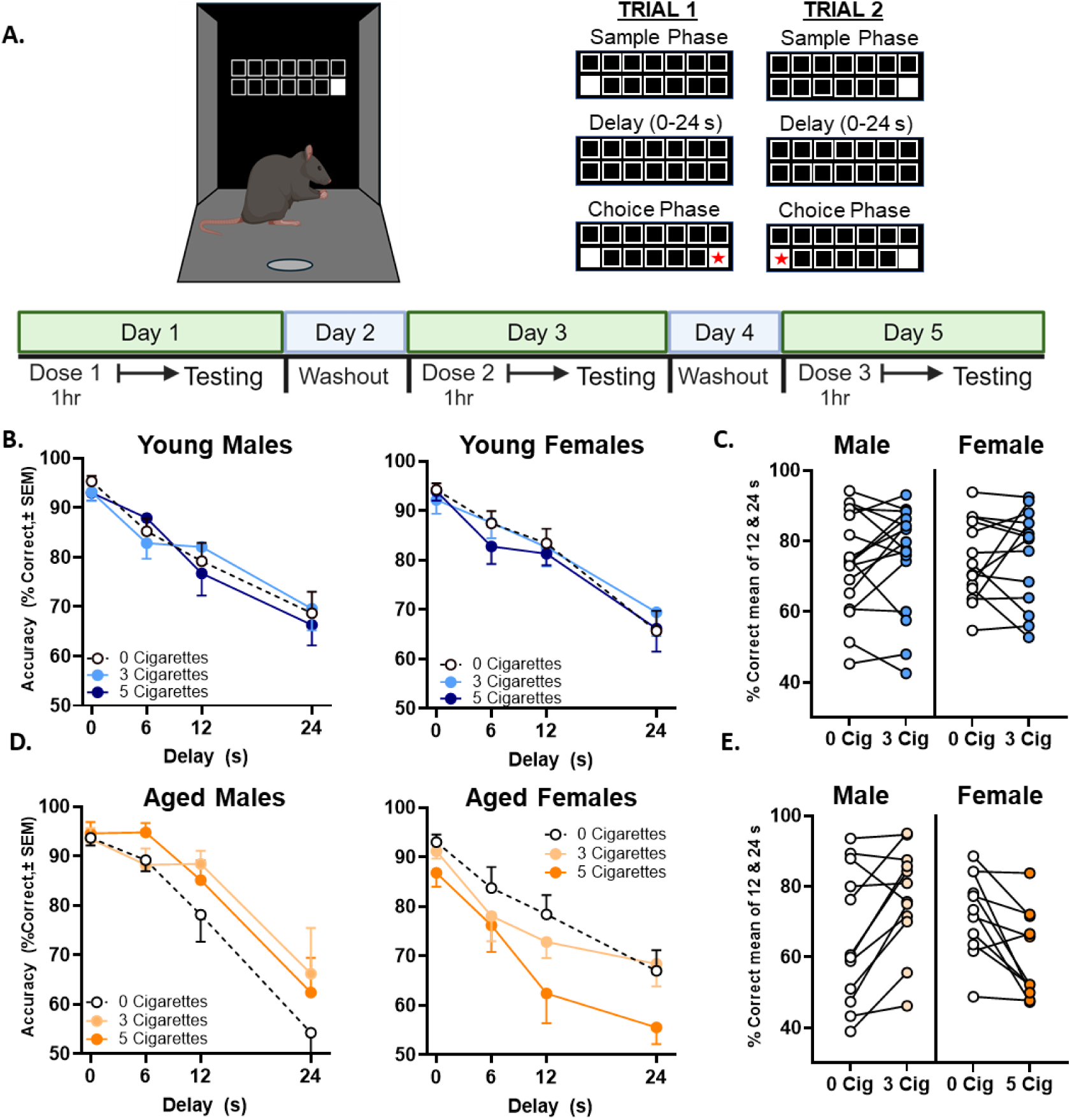
Effects of acute exposure to cannabis smoke on working memory task performance. Young adult (n=17 M, n=14 F) and aged (n=11 M, n=12 F) rats were tested in a non-match-to-sample delayed response task following acute exposure to cannabis smoke, using a within-subjects design. **A.** Top left illustrates the touchscreen apparatus, and top right illustrates two examples of trials in the task (red star indicates correct choice on each trial). Bottom shows timeline of testing with acute smoke exposure. **B.** There was no effect of smoke exposure on working memory accuracy in young adult male or female rats. **C.** There was no significant change in individual performance at 12-24 s delays (averaged) from the 0 to 3 cigarette condition in either young males or females. **D.** Among aged rats, acute cannabis smoke enhanced working memory accuracy in males but impaired accuracy in females. **E.** At the 12-24 s delays (averaged), aged males showed improved accuracy in the 3 cigarette condition, whereas aged females showed impaired accuracy in the 5 cigarette condition. See Table 1 for details of statistical analyses.

A four-factor repeated measures ANOVA (Age x Sex x Cigarettes x Delay) conducted on task accuracy (Figures 1B, D) indicated an Age x Sex x Cigarettes interaction (F(2,100)=4.85, p=.01), such that effects of cannabis smoke exposure on accuracy differed as a function of Age and Sex (see Table 1 for full results). In light of this interaction, the effects of Cigarettes were evaluated in each Age and Sex group. Two-factor ANOVAs (Cigarettes x Delay) conducted separately in aged rats of each sex revealed that in aged males, cannabis smoke *enhanced* working memory accuracy (main effect of Cigarettes, F(2,20)=4.08, p=.03), whereas in aged females, cannabis smoke *impaired* working memory accuracy (main effect of Cigarettes, F(2,22)=8.46, p=.002). In contrast, there were no main effects of interactions involving Cigarettes in young rats of either sex. To better illustrate these results, Figures 1C and E show individual rat accuracy averaged across the longest two delays, at which cannabis produced the largest group effects. The cigarette dose shown is that which produced the largest effect on accuracy in each group (or at the highest dose in young rats).

**Table 1.** Full statistical results for the effects of cannabis smoke exposure on delayed response working memory task accuracy.

| Factor(s) | Result |  |  |  |
| --- | --- | --- | --- | --- |
| Cigs | F (2, 100) =2.308, p=.105 |  |  |  |
| Cigs x Age | F (2, 100) =0.403, p=.669 |  |  |  |
| Cigs x Age x Sex | **F (2, 100) =5.507, p=.005** |  |  |  |
| Delay | **F (2, 100) =4.847, p=.010** |  |  |  |
| Delay x Sex | F (3, 150) =1.378, p=.252 |  |  |  |
| Delay x Age x Sex | F (3, 150) =0.753, p=.522 |  |  |  |
| Cigs x Delay | **F (3, 150) = 3.295, p=.022** |  |  |  |
| Cigs x Delay x Sex | F (6, 300) =1.090, p=.368 |  |  |  |
| Cigs x Delay x Age | F (6, 300) =0.427, p=.861 |  |  |  |
| Cigs x Delay x Age x Sex | F (6, 300) =0.842, p=.538 |  |  |  |
| Age | F (1, 50) =1.439, p=.236 |  |  |  |
| Sex | F (1, 50) =1.601, p=.212 |  |  |  |
| Age x Sex | F (1, 50) =2.280, p=.137 |  |  |  |
|  | Young |  | Aged |  |
| Cigs | F (2, 58) =0.458, p=.635 |  | F (2, 42) =2.361, p=.107 |  |
| Cigs x Sex | F (2, 58) =0.044, p=.957 |  | **F (2, 42) =10.700, p<.001** |  |
| Delay | **F (3, 87) =64.687, p<.001** |  | F (3, 63) =35.385, p <.001 |  |
| Delay x Sex | F (3, 87) =0.456, p=.714 |  | **F (3, 63) =2.794, p=.047** |  |
| Cigs x Delay | F (6, 174) =0.686, p=.661 |  | F (6, 126) =1.446, p=.202 |  |
| Cigs x Delay x Sex | F (6, 174) =1.259, p=.279 |  | F (6, 126) =0.731, p=.625 |  |
| Sex | F (1, 29) =0.036, p=.850 |  | F (1, 21) =3.194, p=.088 |  |
|  | Young |  | Aged |  |
|  | Male | Female | Male | Female |
| Delay | **F(3, 48) =38.499, p<.001** | **F (3, 39) =27.494, p <.001** | **F(3, 30) =16.408, p <.001** | **F (3, 33) =23.348, p <.001** |
| Cigs | F (2, 32) =2.45, p = .784 | F (2, 26) =0.224, p = .801 | **F (2,20) =4.082, p =.033** | **F (2, 22) =8.461, p = .002** |
| Cigs x Delay | F (6, 96) =1.216, p =.305 | F (6, 78) =0.764, p = .600 | F (6, 60) =1.343, p = .252 | F (6, 66) =0.875, p = .518 |

#### Experiment 1b. TUNL task

The TUNL task [38,39] was similar in structure to the working memory task, but no delays were employed. Each trial was divided into sample and choice phases. During the sample phase, one panel (randomly selected from among the 7 positions) was illuminated (Figure 2A). Touching the illuminated panel caused the screen to go blank and the light in the food trough was illuminated. A nosepoke into the food trough caused two panels on the touchscreen to be illuminated (both the original location and a second location separated by 1, 3, or 5 blank locations). A touch on the panel not illuminated during the sample phase (i.e., a “non-match”) yielded a food pellet. A touch on the “incorrect” panel caused the screen to go blank and no food was delivered.

**Figure 2.**
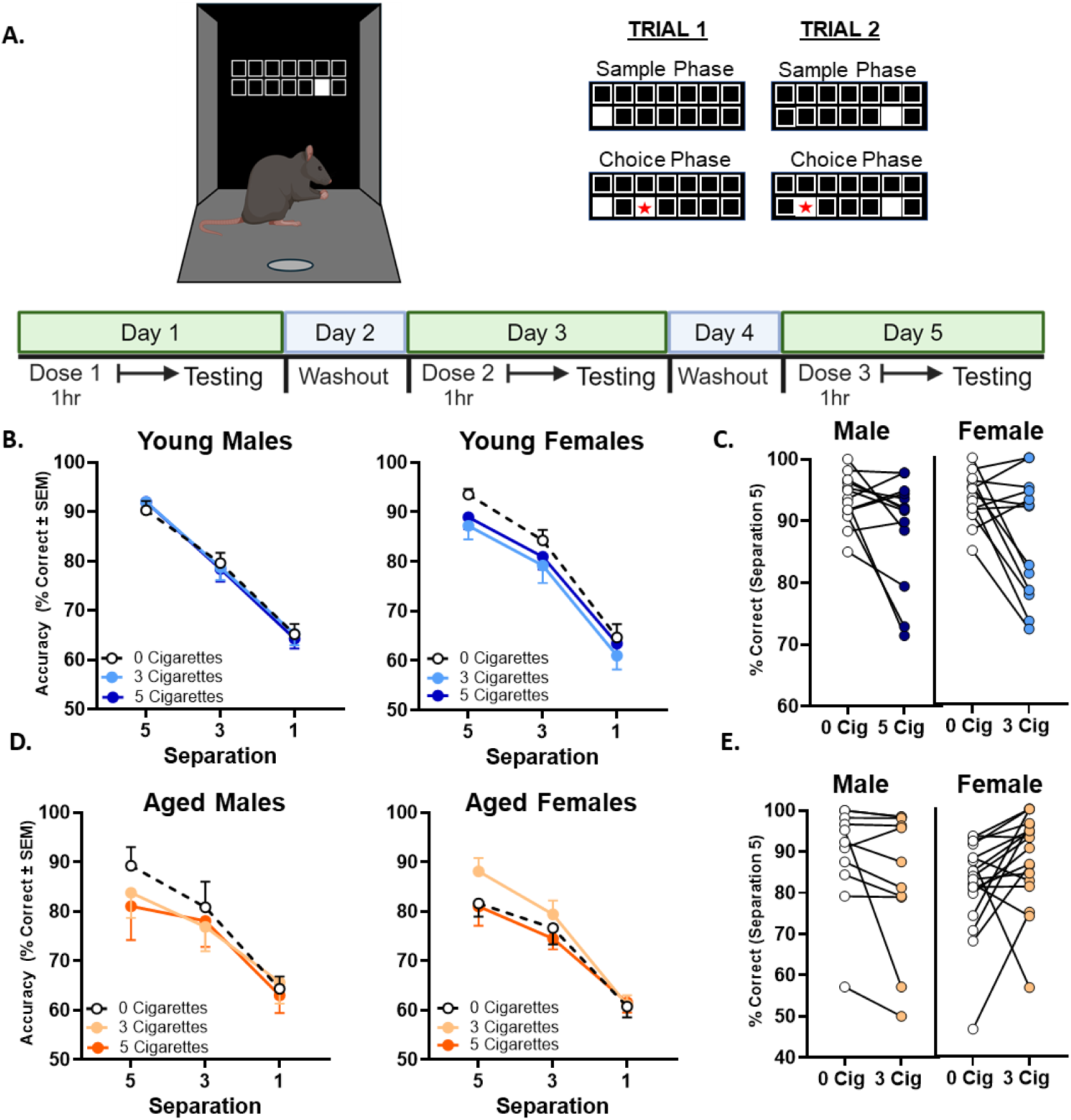
Effects of acute exposure to cannabis smoke on trial-unique non-match-to-location (TUNL) task performance. Young adult (n=17 M, n=13 F) and aged (n=11 M, n=18 F) rats were tested in a trial-unique non-match-to-location task following acute exposure to cannabis smoke, using a within-subjects design. **A.** Top left illustrates the touchscreen apparatus, and top right illustrates two examples of trials in the task (red star indicates correct choice on each trial). Bottom shows timeline of testing with acute smoke exposure. **B-C.** TUNL task accuracy in young adult male and female rats. **D-E.** TUNL task accuracy in aged male and female rats. Although the omnibus ANOVA indicated distinct effects of cannabis smoke in young adult and aged rats of each sex, follow-up analyses revealed no significant effects in any individual age or sex group. See Table 3 for details of statistical analyses.

A four-factor repeated measures ANOVA (Age x Sex x Cigarettes x Separation) revealed an Age x Sex x Cigarettes interaction (F(2,110)=3.77, p=.03), such that effects of cannabis smoke exposure on task accuracy differed as a function of both Age and Sex (Figures 2B, D; see Table 3 for full statistical results). To better elucidate this interaction, two-factor ANOVAs (Cigarettes x Separation) were conducted separately in each Age and Sex group. Unlike in the delayed response task, however, no main effects or interactions involving Cigarettes emerged. To better illustrate these results, Figures 2C and E show individual rat accuracy at the largest separation, at which cannabis produced the greatest group effects. The cigarette dose shown is that which appeared to have the numerically largest effect on accuracy.

**Table 2.**
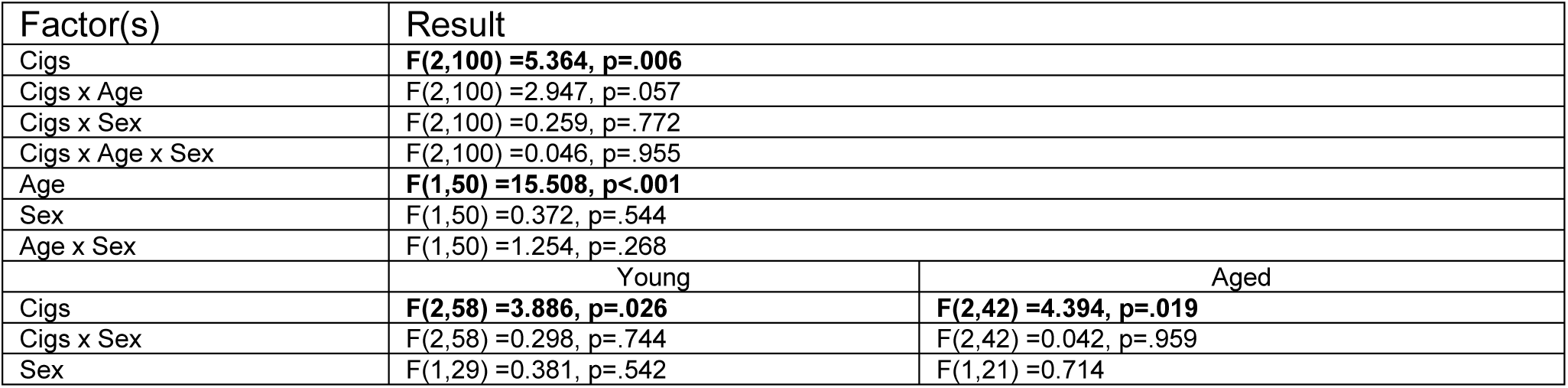
Full statistical results for the effects of cannabis smoke exposure on the number of trials completed per session during the delayed response working memory task.

**Table 3.** Full statistical results for the effects of cannabis smoke exposure on TUNL task accuracy.

| Factor | Result |  |  |  |
| --- | --- | --- | --- | --- |
| Cigs | F(2,110)=1.849, p=.162 |  |  |  |
| Cigs x Age | F (2,110)=1.507, p=.226 |  |  |  |
| Cigs x Age x Sex | **F (2,110)=3.771, p=.026** |  |  |  |
| Separation | **F (2,110) =324.429, p <.001** |  |  |  |
| Separation x Age | **F (2,110)=4.188 p=.018** |  |  |  |
| Separation x Sex | **F (2,110)=1.181 p=.311** |  |  |  |
| Separation x Age x Sex | F (2,110)=0.782 p=.460 |  |  |  |
| Cigs x Separation | F(4,220)=0.423 p=.792 |  |  |  |
| Cigs x Separation x Sex | F (4,220)=0.321, p=.863 |  |  |  |
| Cigs x Separation x Age | F (4,220)=0.538, p=.708 |  |  |  |
| Cigs x Separation x Age x Sex | F(4,220)=1.272, p=.282 |  |  |  |
| Age | F (1,55) =2.632, p=.110 |  |  |  |
| Sex | F (1,55) =0.295, p=.589 |  |  |  |
| Age x Sex | F (1,55) =0.172, p=.680 |  |  |  |
|  | Young |  | Aged |  |
| Cigs | F (2,56)=1.635, p=.204 |  | F(2,54)= 1.689, p=.194 |  |
| Cigs x Sex | F(2,56)=1.989, p=.146 |  | F (2,54) =1.802, p=.175 |  |
| Separation | **F (2,56)= 336.821, p<.001** |  | **F(2,54)=90.031, p<.001** |  |
| Separation x Sex | F(2,56)=2.817, p=.068 |  | F(2,54)=0.214, p=.808 |  |
| Cigs x Separation | F (4,112)=0.071, p=.991 |  | F(4,108)=0.540, p=.707 |  |
| Cigs x Separation x Sex | F (4,112)=0.399, p=.809 |  | F(4,108)=1.181, p=.323 |  |
| Sex | F(1,29) =0.016, p=.899 |  | F(1,27) =0.295, p=.591 |  |
|  | Young |  | Aged |  |
|  | Male | Female | Male | Female |
| Separation | **F(2,32)=190.218, p<.001** | **F (2,24)=153.880, p<.001** | **F(2,20)=30.260, p<.001** | **F (2,34)=68.675, p<.001** |
| Cigs | F (2,32)=0.035, p=.966 | F (2,24)=2.626, p=.093 | F (2,20)=1.452, p=.258 | F (2,34)=2.243, p=.122 |
| Cigs x Sep | F (4,64)=0.228, p=.922 | F (4,48)=0.269, p=.896 | F (4,40)=56.673, p=.701 | F (4,68)=1.331, p=.267 |

### Experiment 2: Effects of repeated oral Δ9THC on cognition in young adult and aged rats

Oral consumption is one of the most prevalent routes of cannabis use in older adults [43]. Experiment 2 was designed to mimic this pattern of consumption to determine its effects on cognition. Rats were trained on a delayed response working memory task until stable performance emerged and then were divided into two groups matched for performance. Both groups were initially trained to consume plain gelatin (to overcome neophobia), followed by daily sessions in which they were given 1 h home cage access to 10.0 g of plain gelatin (the control group) or gelatin containing Δ9THC (1.0 mg/kg, the highest dose that rats would reliably consume on a daily basis; the THC group). Rats consumed either Δ9THC or control gelatin daily, while continuing behavioral testing. After 3 weeks of testing in the working memory task, rats were assessed in the Morris water maze under the same conditions. Gelatin consumption took place in the afternoons and behavioral testing was conducted in the mornings, to eliminate potential acute effects of Δ9THC on performance.

#### Delayed match-to-sample working memory task

The delayed response task was similar to that used in Experiment 1, but was conducted in standard operant chambers equipped with two retractable levers and a food delivery trough. The task consisted of multiple trials in 40-min sessions. Each trial began with a single lever (the sample, either left or right) extended into the chamber (Figure 4A). A press initiated the delay phase (0-24 s), during which rats nosepoked into the food trough. This was followed by the choice phase, in which both levers extended. A press on the same lever extended in the sample phase (“correct”) yielded a food pellet (i.e., rats had to employ a “match” rule). A press on the opposite (“incorrect”) lever caused levers to retract and no food delivery.

**Figure 3.**
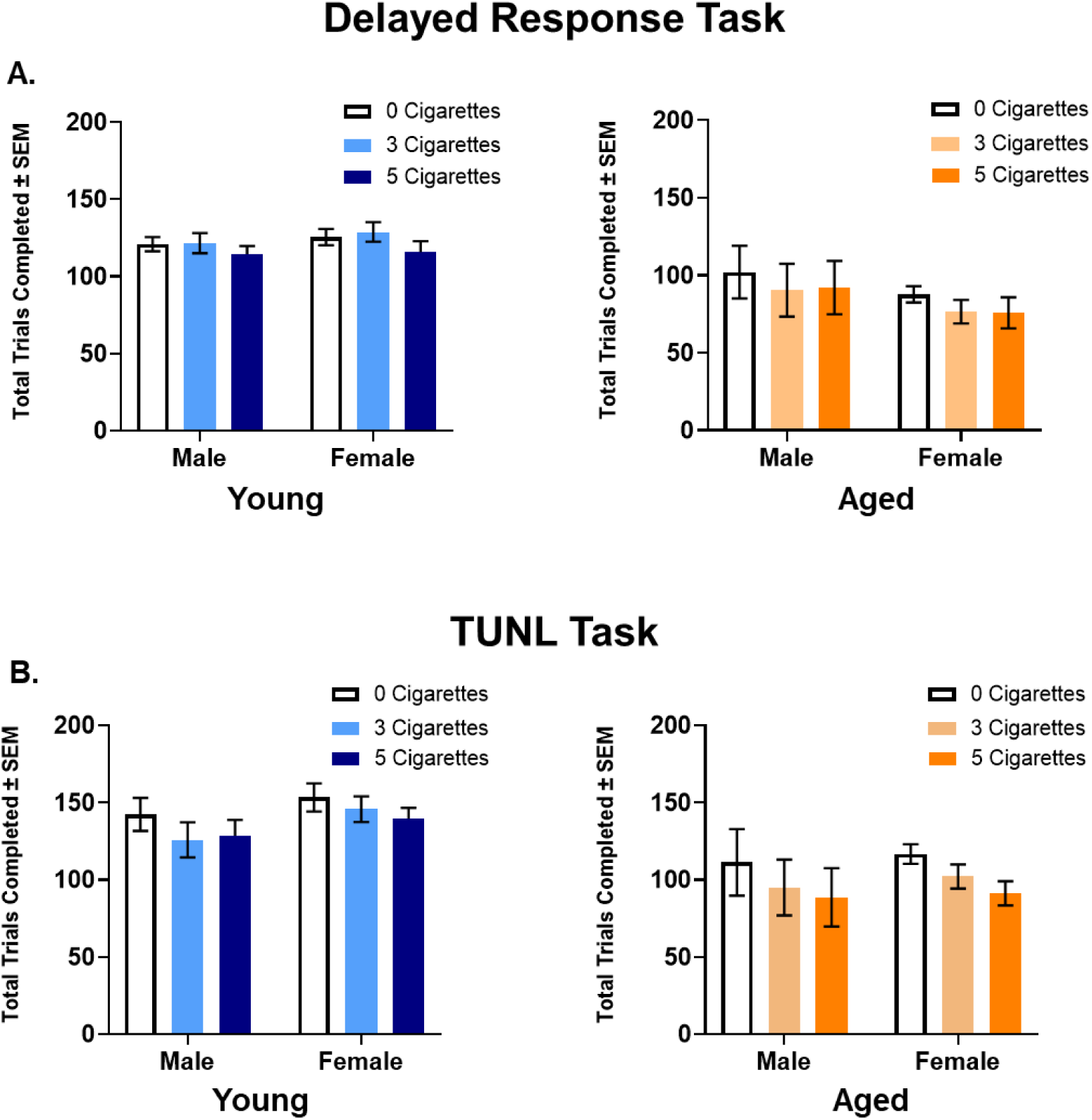
Effects of acute exposure to cannabis smoke on the number of trials completed per session in the delayed response and TUNL tasks (Experiment 1). **A.** The number of trials completed in young adult and aged rats on the non-match-to-sample delayed response task following acute exposure to cannabis smoke. **B.** The number of trials completed in young adult and aged rats on the TUNL task following acute exposure to cannabis smoke. See Tables 2 and 4 for full statistical results.

**Figure 4.**
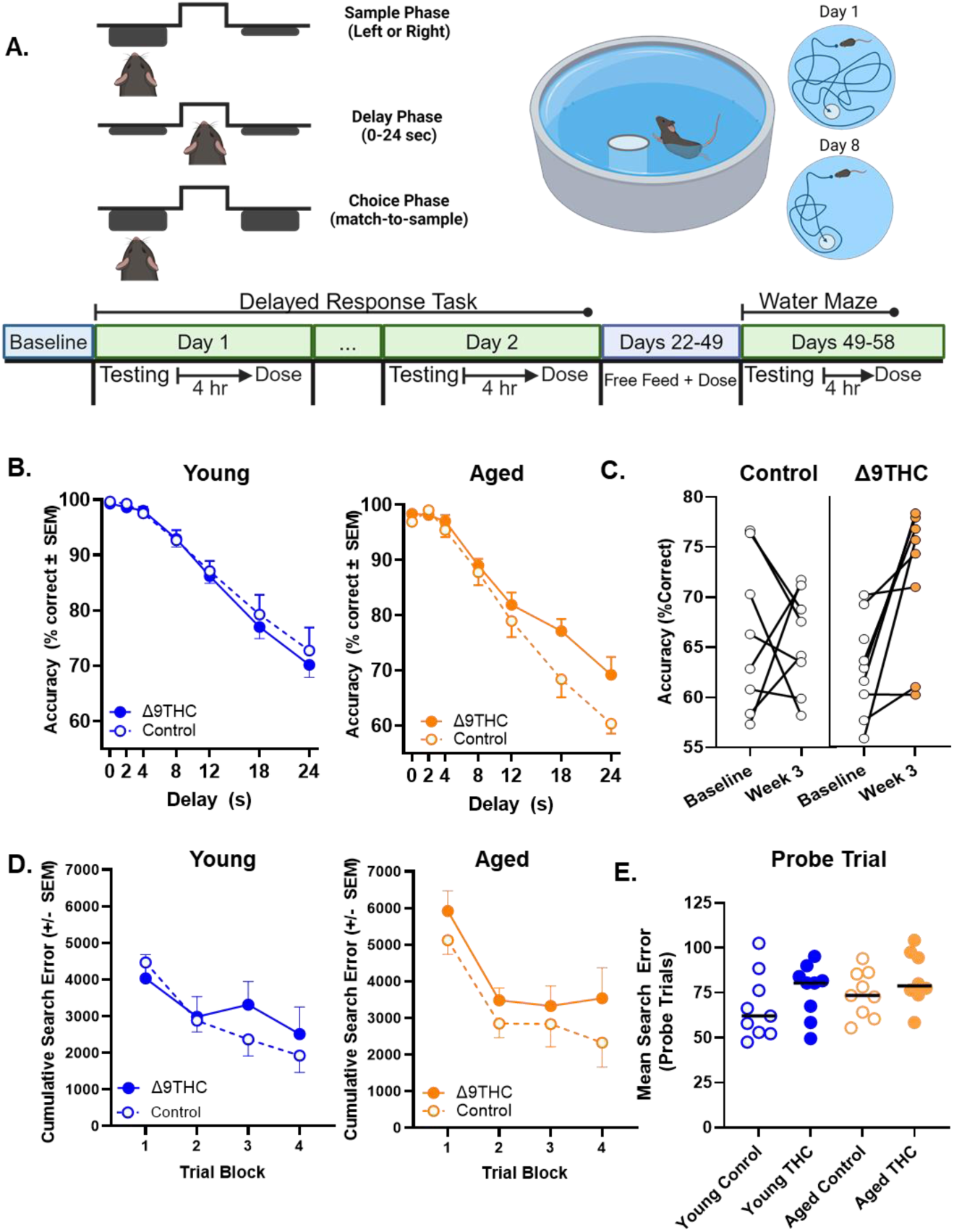
Effects of chronic oral consumption of Δ9THC on working and spatial memory. Young adult (n=10 control (5M, 5F), n=10 Δ9THC (5M, 5F)) and aged (n=9 control (5M, 4F), n=9 Δ9THC (5M, 4F)) rats were tested in a match-to-sample delayed response task and the Morris water maze task, while undergoing daily consumption of Δ9THC or plain (control) gelatin. **A.** Top left illustrates a single trial in the working memory task, and top right illustrates the water maze apparatus and swim paths on two example trials. Bottom shows timeline of testing with chronic Δ9THC consumption. **B.** At 3 weeks of gelatin consumption, there was no effect of Δ9THC on working memory accuracy in young adult rats. In contrast, Δ9THC enhanced working memory accuracy in aged rats, particularly at long delays. **C.** Among aged rats, those in the control group showed no change in accuracy from baseline (pre-gelatin) to Week 3, whereas those in the Δ9THC group showed a significant improvement in accuracy across the same time period. **D.** In the water maze task, there were no significant effects of Δ9THC on search accuracy (cumulative search error) in either age group during training trials. **E.** Similarly, there were no significant effects of Δ9THC on memory for the platform location during the interpolated probe trials (mean search error, averaged across all four trials). Note that as there were no main effects or interactions involving sex, data are collapsed across sex for all measures. See Tables 5 and 9 for details of statistical analyses.

Prior to Δ9THC or control gelatin consumption, comparison of working memory accuracy (four-factor ANOVA, Age x Drug x Sex x Delay) revealed no main effects or interactions involving Drug (Fs<1.65, ps>.13), indicating comparable performance between drug groups prior to Δ9THC consumption (Figure 5A and Table 6). Rats regularly consumed all the gelatin during consumption sessions, with no differences across groups (Figure 5B and Table 7). To evaluate effects of chronic Δ9THC on working memory, data were averaged across the last five sessions of testing in Week 3 and assessed using a four-factor ANOVA (Age x Drug x Sex x Delay). There was a main effect of Age (F(1,180)=10.38, p=.003) and a Delay x Age interaction (F(6,180)=2.88, p=.01), such that aged rats performed worse than young, particularly at longer delays (Figure 4B). Most importantly, there was a Delay x Age x Drug interaction (F(6,180)=2.45, p=.03), indicating that, across delays, Δ9THC had distinct effects in young and aged rats.

**Figure 5.**
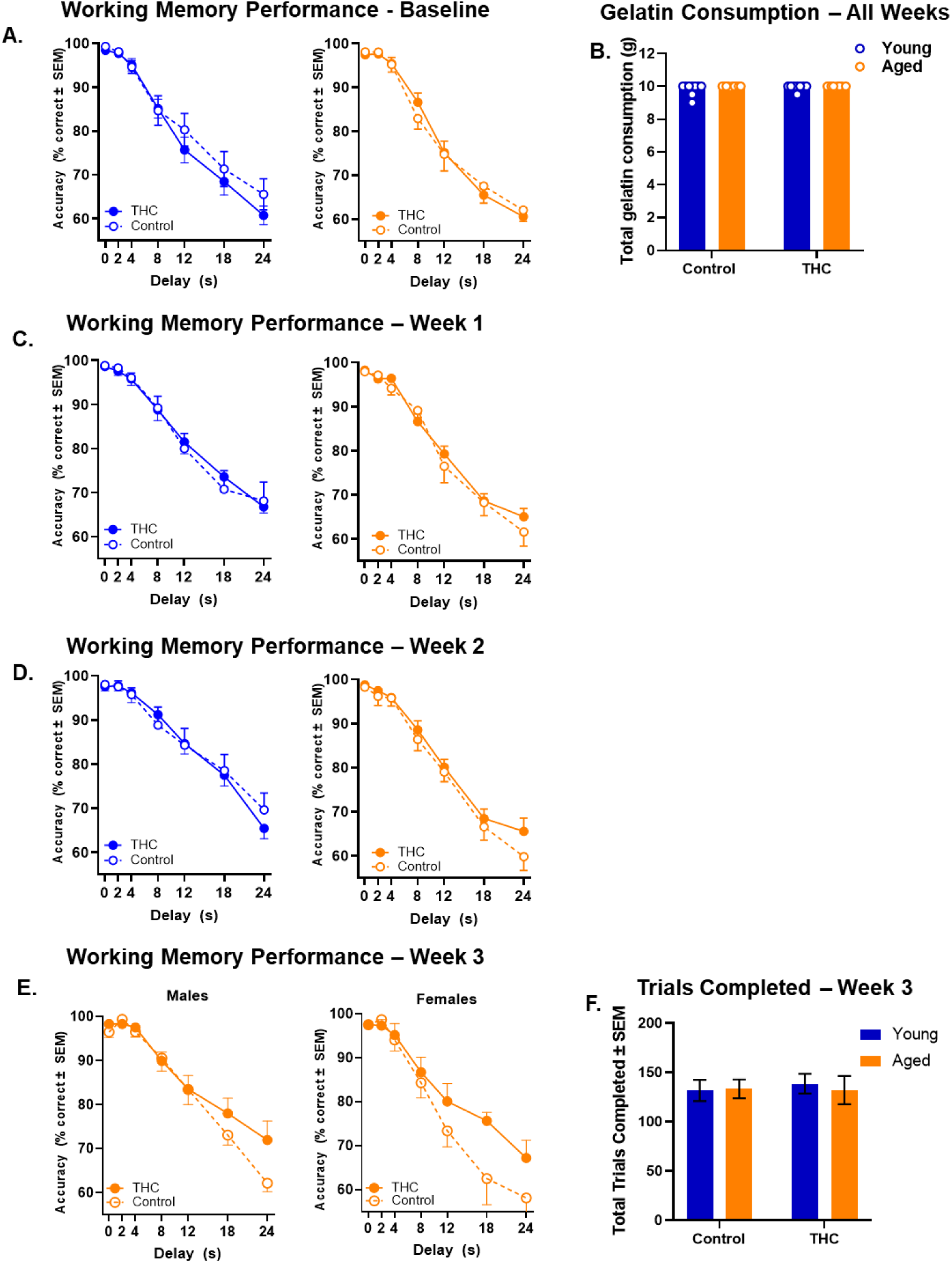
Effects of chronic oral Δ9THC consumption on working memory in young adult and aged rats (Experiment 2). **A.** During baseline performance (prior to gelatin consumption), there were no significant effects or interactions involving age, sex, or drug group assignment. **B.** There were no significant age or sex differences in the quantity of Δ9THC vs. control gelatin consumed. **C.** There were no significant main effects or interactions involving Δ9THC condition on working memory performance during Week 1 of gelatin consumption. **D.** There were no significant main effects or interactions involving Δ9THC condition on working memory performance during Week 2 of gelatin consumption. **E.** Week 3 performance for aged animals split by sex. **F.** There were no significant differences between age or drug condition in the number of trials completed on the working memory task during Week 3 of gelatin consumption (averaged across all sessions). See Tables 6-8 for full statistical results

To further explore this interaction, three-factor ANOVAs (Delay x Sex x Drug) were conducted separately in each age group. Among aged rats, there was a significant Delay x Drug interaction (F(6,84)=2.76, p=.02), indicating enhanced accuracy in the Δ9THC compared to the control group, particularly at longer delays (Figure 4B). In contrast, the same analysis in young rats revealed no main effect or interaction involving Drug (see Table 5 for full results). To better assess Δ9THC effects in aged rats, Fig. 3C shows averaged accuracy across the longest two delays at baseline prior to drug consumption and at the 3-week timepoint. Whereas aged control rats exhibited no change in performance across weeks (t(16)=0.12, p=.90), aged rats consuming Δ9THC showed a significant improvement across this period (t(16)=3.31), p=.004), supporting the interpretation that chronic oral Δ9THC alleviates age-associated working memory impairment.

**Table 4.** Full statistical results for the effects of cannabis smoke exposure on the number of trials completed per session during the TUNL task.

| Factor | Result |  |  |
| --- | --- | --- | --- |
| Cigs | **F(2,110) = 11.967, p<.001** |  |  |
| Cigs x Sex | F(2,110) = 0.394, p=.675 |  |  |
| Cigs x Age | F(2, 110) = 0.850, p=.430 |  |  |
| Cigs x Age x Sex | F(2,110) = 0.099, p=.906 |  |  |
| Age | **F(1,55) = 13.711, p<.001** |  |  |
| Sex | F(1,55) = 0.836, p=.365 |  |  |
| Age x Sex | F(1,55) = 0.182, p= .671 |  |  |
|  | Young |  | Aged |
| Cigs | F(2,56) = 2.538, p=.088 |  | **F(2,54) = 16.881, p<.001** |
| Cigs x Sex | F(2, 56) = 0.290, p=.749 |  | F(2,54) = 0.159, p=.854 |
| Sex | F(1,28) = 1.343, p=.256 |  | F(1,27) = 0.087, p=.770 |

**Table 5.** Full statistical results for the effects of chronic Δ9THC consumption on delayed response working memory task accuracy (in Week 3)

| Factor | Result |  |
| --- | --- | --- |
| Delay | **F(6,180) = 240.685, p<.001** |  |
| Delay x Sex | F(6,180)=0.335, p=.918 |  |
| Delay x Drug | F(6,180)=0.881, p=.510 |  |
| Delay x Age | **F(6,180)= 2.882, p=.010** |  |
| Delay x Sex x Drug | F(1,180)=0.395, p=.882 |  |
| Delay x Sex x Age | **F(6,180)=3.140, p=.006** |  |
| Delay x Drug x Age | **F(6,180)=2.451, p=.027** |  |
| Delay x Sex x Age x Drug | F(6,180)=0.581, p=.745 |  |
| Age | **F(1,30)=10.377, p=.003** |  |
| Sex | F(1,30)=0.000, p=.986 |  |
| Drug | F(1,30)=1.212, p=.280 |  |
| Age x Sex | **F(1,30)=7.953, p=.008** |  |
| Age x Group | F(1,30)=3.239, p=.082 |  |
| Sex x Group | F(1,30)=0.418, p=.523 |  |
| Age x Sex x Group | F(1,30)=0.160, p=.692 |  |
|  | Young | Aged |
| Delay | **F(6,96)=108.546, p<.001** | **F(6,84)=131.401, p<.001** |
| Delay x Sex | F(6,96)= 1.949, p=.081 | F(6,84)=1.548, p=.173 |
| Delay x Drug | F(6,96)=0.302, p=.935 | **F(6,84)=2.763, p=.017** |
| Delay x Sex x Drug | F(6,96)=0.207, p=.974 | F(6,84)=0.713, p=.640 |
| Sex | F(1,16)=3.622, p=.075 | F(1,14)=4.677, p=.048 |
| Drug | F(1,16)=0.220, p=.646 | F(1,14)=5.009, p=.042 |
| Sex x Drug | F(1,16)=0.027, p=.871 | F(1,14)=0.651, p=.433 |

**Table 6.**
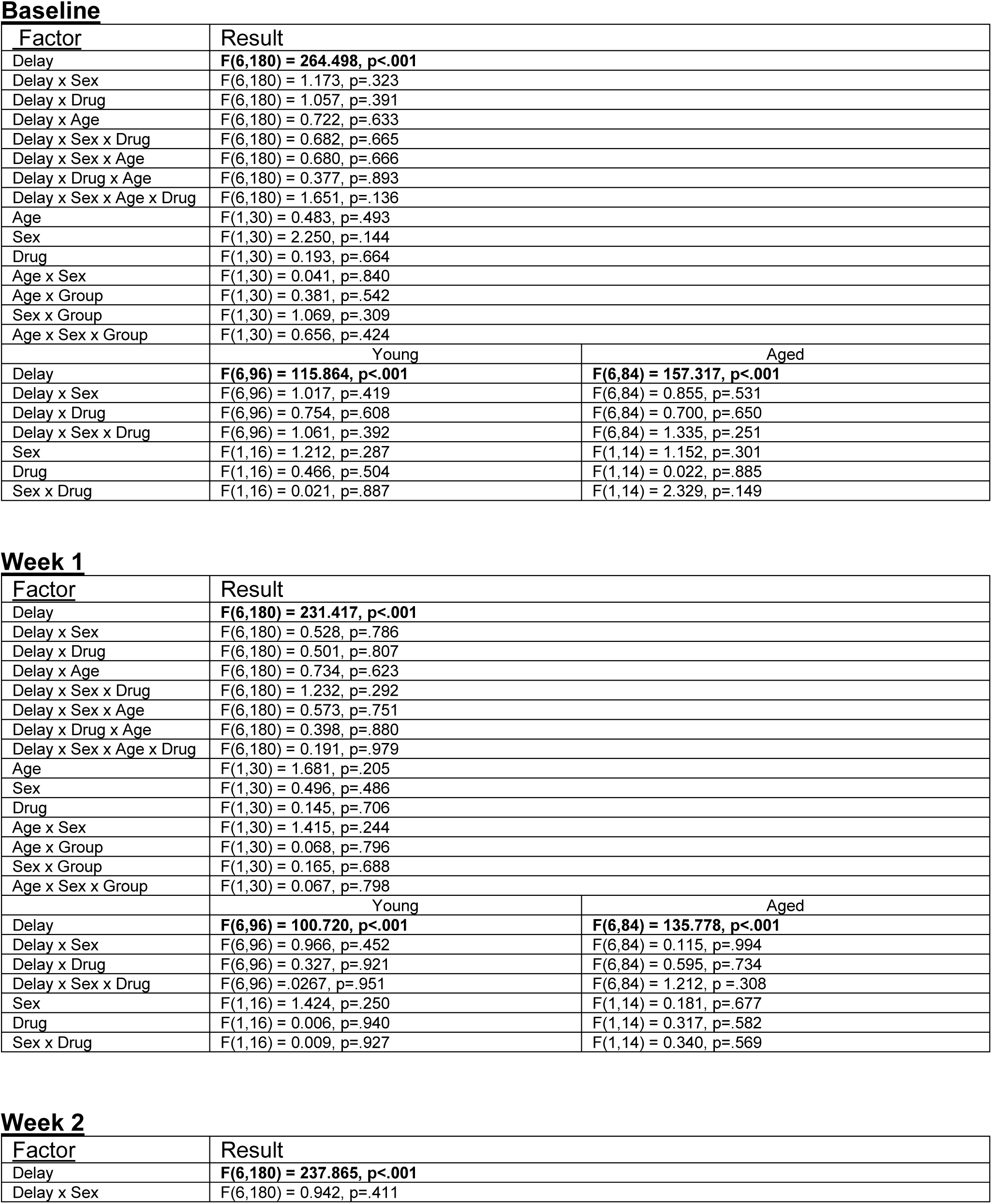
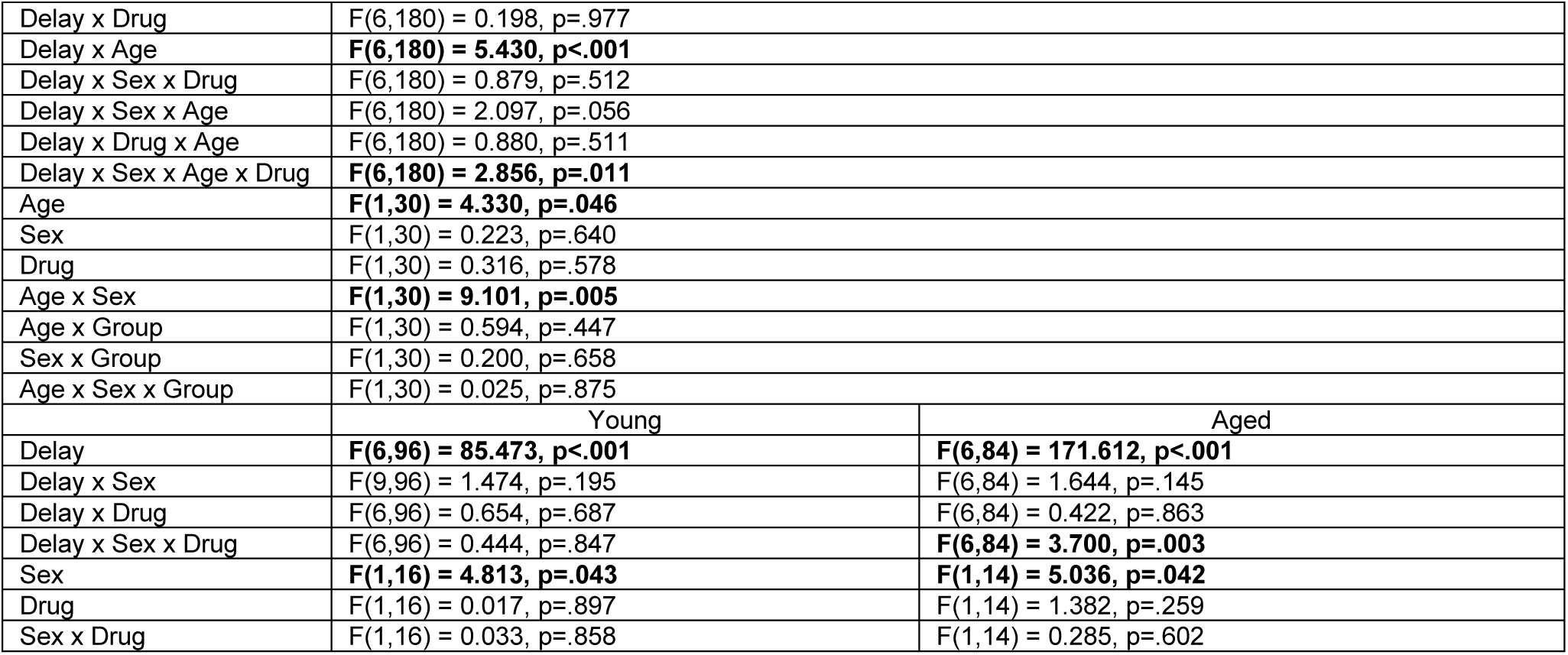
Full statistical results for the effects of chronic Δ9THC consumption on delayed response working memory task accuracy during baseline (prior to gelatin consumption) and during Weeks 1 and 2.

**Baseline**
| Factor | Result |  |
| --- | --- | --- |
| Delay | **F(6,180) = 264.498, p<.001** |  |
| Delay x Sex | F(6,180) = 1.173, p=.323 |  |
| Delay x Drug | F(6,180) = 1.057, p=.391 |  |
| Delay x Age | F(6,180) = 0.722, p=.633 |  |
| Delay x Sex x Drug | F(6,180) = 0.682, p=.665 |  |
| Delay x Sex x Age | F(6,180) = 0.680, p=.666 |  |
| Delay x Drug x Age | F(6,180) = 0.377, p=.893 |  |
| Delay x Sex x Age x Drug | F(6,180) = 1.651, p=.136 |  |
| Age | F(1,30) = 0.483, p=.493 |  |
| Sex | F(1,30) = 2.250, p=.144 |  |
| Drug | F(1,30) = 0.193, p=.664 |  |
| Age x Sex | F(1,30) = 0.041, p=.840 |  |
| Age x Group | F(1,30) = 0.381, p=.542 |  |
| Sex x Group | F(1,30) = 1.069, p=.309 |  |
| Age x Sex x Group | F(1,30) = 0.656, p=.424 |  |
|  | Young | Aged |
| Delay | **F(6,96) = 115.864, p<.001** | **F(6,84) = 157.317, p<.001** |
| Delay x Sex | F(6,96) = 1.017, p=.419 | F(6,84) = 0.855, p=.531 |
| Delay x Drug | F(6,96) = 0.754, p=.608 | F(6,84) = 0.700, p=.650 |
| Delay x Sex x Drug | F(6,96) = 1.061, p=.392 | F(6,84) = 1.335, p=.251 |
| Sex | F(1,16) = 1.212, p=.287 | F(1,14) = 1.152, p=.301 |
| Drug | F(1,16) = 0.466, p=.504 | F(1,14) = 0.022, p=.885 |
| Sex x Drug | F(1,16) = 0.021, p=.887 | F(1,14) = 2.329, p=.149 |

**Week 1**
| Factor | Result |  |
| --- | --- | --- |
| Delay | **F(6,180) = 231.417, p<.001** |  |
| Delay x Sex | F(6,180) = 0.528, p=.786 |  |
| Delay x Drug | F(6,180) = 0.501, p=.807 |  |
| Delay x Age | F(6,180) = 0.734, p=.623 |  |
| Delay x Sex x Drug | F(6,180) = 1.232, p=.292 |  |
| Delay x Sex x Age | F(6,180) = 0.573, p=.751 |  |
| Delay x Drug x Age | F(6,180) = 0.398, p=.880 |  |
| Delay x Sex x Age x Drug | F(6,180) = 0.191, p=.979 |  |
| Age | F(1,30) = 1.681, p=.205 |  |
| Sex | F(1,30) = 0.496, p=.486 |  |
| Drug | F(1,30) = 0.145, p=.706 |  |
| Age x Sex | F(1,30) = 1.415, p=.244 |  |
| Age x Group | F(1,30) = 0.068, p=.796 |  |
| Sex x Group | F(1,30) = 0.165, p=.688 |  |
| Age x Sex x Group | F(1,30) = 0.067, p=.798 |  |
|  | Young | Aged |
| Delay | **F(6,96) = 100.720, p<.001** | **F(6,84) = 135.778, p<.001** |
| Delay x Sex | F(6,96) = 0.966, p=.452 | F(6,84) = 0.115, p=.994 |
| Delay x Drug | F(6,96) = 0.327, p=.921 | F(6,84) = 0.595, p=.734 |
| Delay x Sex x Drug | F(6,96) = 0.0267, p=.951 | F(6,84) = 1.212, p=.308 |
| Sex | F(1,16) = 1.424, p=.250 | F(1,14) = 0.181, p=.677 |
| Drug | F(1,16) = 0.006, p=.940 | F(1,14) = 0.317, p=.582 |
| Sex x Drug | F(1,16) = 0.009, p=.927 | F(1,14) = 0.340, p=.569 |

**Week 2**
| Factor | Result |
| --- | --- |
| Delay | **F(6,180) = 237.865, p<.001** |
| Delay x Sex | F(6,180) = 0.942, p=.411 |
| Delay x Drug | F(6,180) = 0.198, p=.977 |  |
| Delay x Age | **F(6,180) = 5.430, p<.001** |  |
| Delay x Sex x Drug | F(6,180) = 0.879, p=.512 |  |
| Delay x Sex x Age | F(6,180) = 2.097, p=.056 |  |
| Delay x Drug x Age | F(6,180) = 0.880, p=.511 |  |
| Delay x Sex x Age x Drug | **F(6,180) = 2.856, p=.011** |  |
| Age | **F(1,30) = 4.330, p=.046** |  |
| Sex | F(1,30) = 0.223, p=.640 |  |
| Drug | F(1,30) = 0.316, p=.578 |  |
| Age x Sex | **F(1,30) = 9.101, p=.005** |  |
| Age x Group | F(1,30) = 0.594, p=.447 |  |
| Sex x Group | F(1,30) = 0.200, p=.658 |  |
| Age x Sex x Group | F(1,30) = 0.025, p=.875 |  |
|  | Young | Aged |
| Delay | **F(6,96) = 85.473, p<.001** | **F(6,84) = 171.612, p<.001** |
| Delay x Sex | F(9,96) = 1.474, p=.195 | F(6,84) = 1.644, p=.145 |
| Delay x Drug | F(6,96) = 0.654, p=.687 | F(6,84) = 0.422, p=.863 |
| Delay x Sex x Drug | F(6,96) = 0.444, p=.847 | **F(6,84) = 3.700, p=.003** |
| Sex | **F(1,16) = 4.813, p=.043** | **F(1,14) = 5.036, p=.042** |
| Drug | F(1,16) = 0.017, p=.897 | F(1,14) = 1.382, p=.259 |
| Sex x Drug | F(1,16) = 0.033, p=.858 | F(1,14) = 0.285, p=.602 |

**Table 7.**
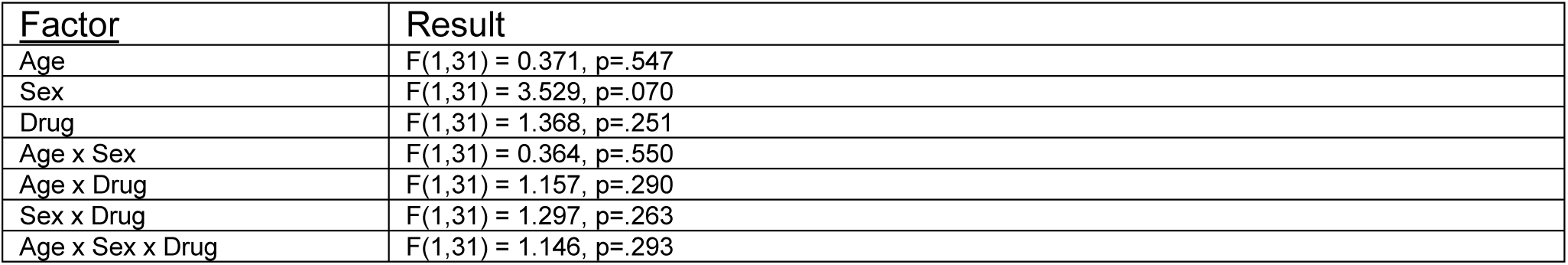
Full statistical results for the effects of Age, Sex, and Δ9THC on gelatin consumption during testing in the delayed response working memory task.

**Table 8.** Full statistical results for the effects of Δ9THC consumption on the number of trials completed per session during the delayed response working memory task in Week 3.

| Factor | Result |
| --- | --- |
| Age | $F(1,31) = 0.526, p=.474$ |
| Sex | $F(1,31) = 1.520, p=.227$ |
| Drug | $F(1,31) = 0.680, p=.416$ |
| Age x Sex | $F(1,31) = 1.081, p=.307$ |
| Age x Drug | $F(1,31) = 1.070, p=.309$ |
| Sex x Drug | $F(1,31) = 0.884, p=.354$ |
| Age x Sex x Drug | $F(1,31) = 1.199, p=.282$ |

#### Morris water maze task

The maze consisted of a tank filled with opaque water in which an escape platform was hidden beneath the surface in one quadrant (spatial task, Figure 4A) or protruded above the surface and moved locations on each trial (cued task) [18,44,45]. To assess spatial memory, rats received three trials/day over eight days. On each training trial, rats were started at a different location and swam until finding the hidden platform. Every sixth trial was a 30-s probe trial on which the platform was made unavailable for escape. After eight days of spatial training, rats were assessed on the cued task to evaluate visual acuity, sensorimotor abilities, and motivation to escape the water.

A four-factor repeated measures ANOVA (Age x Sex x Drug x Trial Block) conducted on performance across training trials (Figure 4D) revealed a main effect of Trial Block, such that rats improved their performance (reduced search error) across training (F(3,78)=18.05 p<.001), as well as a trend toward a main effect of Age (F(1,29)=4.00 p=.056) such that aged rats had less accurate searches than young, but no main effects or interactions involving Drug. A similar pattern of results was evident on probe trials (Figure 4E), as well as on pathlength data on the cued task (see Table 9 and Figure 6 for full results), indicating that chronic Δ9THC consumption did not affect either spatial or cued task performance.

**Table 9.** Full statistical results for the effects of chronic Δ9THC consumption on water maze performance (both cued and spatial task versions)

| Factor | Result |
| --- | --- |
| **Cue Training – Pathlength (trials 2-6)** |  |
| Age | **F(1,27) = 9.971, p=.004** |
| Sex | F(1,27) = 0.182, p=.673 |
| Drug | F(1,27) = 0.015, p=.903 |
| Age x Sex | F(1,27) = 0.001, p=.974 |
| Age x Drug | F(1,27) = 0.159, p=.693 |
| Sex x Drug | F(1,27) = 2.270, p=.144 |
| Age x Sex x Drug | F(1,27) = 0.273, p=.605 |
| **Training Trials - Cumulative Search Error** |  |
| Block | **F(3,81) = 20.498, p<.001** |
| Block x Age | F(3,81) = 1.277, p=.288 |
| Block x Sex | **F(3,81) = 3.269, p=.025** |
| Block x Drug | F(3,81) = 0.498, p=.685 |
| Block x Age x Sex | F(3,81) = 1.583, p=.200 |
| Block x Age x Drug | F(3,81) = 0.650, p=.585 |
| Block x Sex x Drug | F(3,81) = 0.412, p=.745 |
| Block x Age x Sex x Drug | F(3,81) = 0.304, p=.823 |
| Age | F(1,27) = 2.950, p=.097 |
| Sex | **F(1,27) = 7.338, p=.012** |
| Drug | F(1,27) = 3.006, p=.094 |
| Age x Sex | F(1,27) = 2.192, p=.150 |
| Age x Drug | F(1,27) = 0.173, p=.681 |
| Sex x Drug | F(1,27) = 0.234, p=.632 |
| Age x Sex x Drug | F(1,27) = 0.300, p=.588 |
| **Probe Trials – Mean Search Error** |  |
| Probe | **F(3,81) = 12.715, p<.001** |
| Probe x Age | F(3,81) = 0.160, p=.923 |
| Probe x Sex | F(3,81) = 0.367, p=.777 |
| Probe x Drug | F(3,81) = 0.919, p=.435 |
| Probe x Age x Sex | F(3,81) = 0.752, p=.524 |
| Probe x Age x Drug | F(3,81) = 0.299, p=.826 |
| Probe x Sex x Drug | F(3,81) = 0.361, p=.782 |
| Probe x Age x Sex x Drug | F(3,81) = 1.449, p=.235 |
| Age | F(1,27) = 1.628, p=.213 |
| Sex | F(1,27) = 1.640, p=.211 |
| Drug | F(1,27) = 2.674, p=.114 |
| Age x Sex | F(1,27) = 0.043, p=.838 |
| Age x Drug | F(1,27) = 0.037, p=.849 |
| Sex x Drug | F(1,27) = 0.011, p=.917 |
| Age x Sex x Drug | F(1,27) = 0.137, p=.681 |

**Figure 6.**
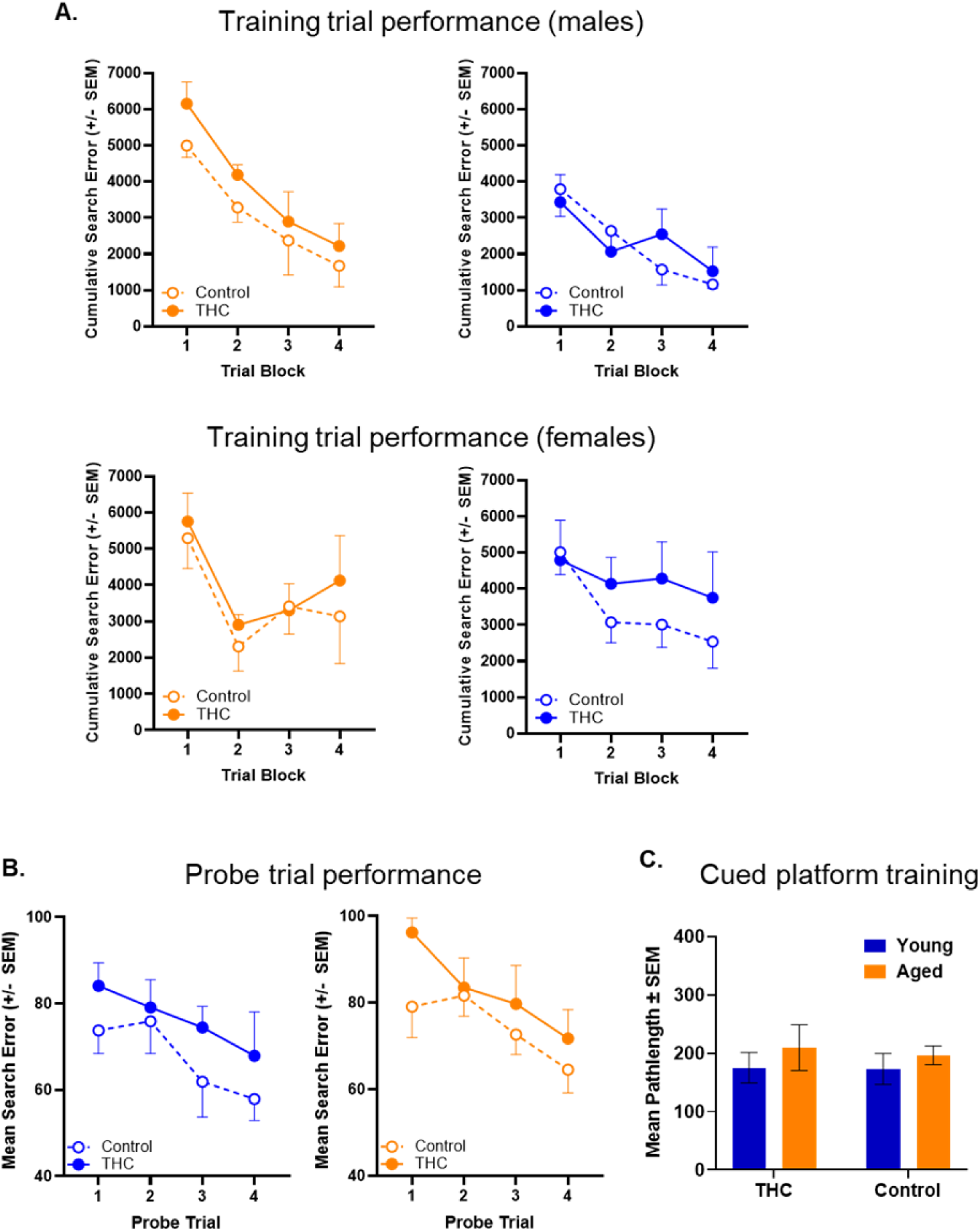
Effects of chronic oral Δ9THC consumption on water maze performance in young adult and aged rats (Experiment 2). **A.** Cumulative search errors across blocks of training trials in the hidden platform (spatial) task, separated by sex. There were no significant main effects or interactions involving Δ9THC condition. **B.** Performance (mean search errors) across the four interpolated probe trials. There were no significant main effects or interactions involving Δ9THC condition. **C.** Performance (pathlength to reach the platform, averaged across trials 2-6) on the cued (visible platform) version of the water maze task. There were no significant main effects or interactions involving Δ9THC condition. See Table 9 for full statistical results.

### Experiment 3: Pharmacokinetics of Δ9THC and its metabolites after different routes of administration

To evaluate Δ9THC pharmacokinetics, rats were exposed to smoke from 5 cannabis cigarettes or consumed gelatin containing 1.0 mg/kg Δ9THC as in Experiments 1 and 2. Blood was collected via tail nicks at different timepoints after consumption and evaluated for Δ9THC and two major metabolites (11-OH-Δ9THC and 11-COOH-Δ9THC) via ultraperformance liquid chromatography-tandem mass spectrometry (UPLC-MS/MS) as in [46,47]. Plasma concentrations of analytes for which sufficient numbers of samples were above lower limit of quantification (LLOQ, 2.5 ng/mL) were analyzed in SPSS.

Following cannabis smoke exposure, peak plasma Δ9THC concentrations in both young adult and aged rats were achieved 10 min after smoke exposure (Figure 7A). Although aged rats tended to have higher C_max_ values than young, these differences were not reliable (two-factor ANOVA, main effect of Age, F(1,33)=3.34, p=.08; Age x Sex interaction, F(1,33)=0.120, p=.73). Comparison of plasma Δ9THC at 10-60 min (the timepoints at which there were sufficient data to analyze; Figure 7A) using a three-factor ANOVA (Age x Sex x Timepoint) also revealed no main effect of Age (F(1,33)=0.95, p=.34), or Age x Timepoint interaction (F(3,99)=2.40, p=.07; see Table 10 for full results). There were too few data points to evaluate the short-lived 11-OH-Δ9THC metabolite (Table 11); however, analysis of the secondary metabolite 11-COOH-Δ9THC revealed no group differences (Figure 7B).

**Figure 7.**
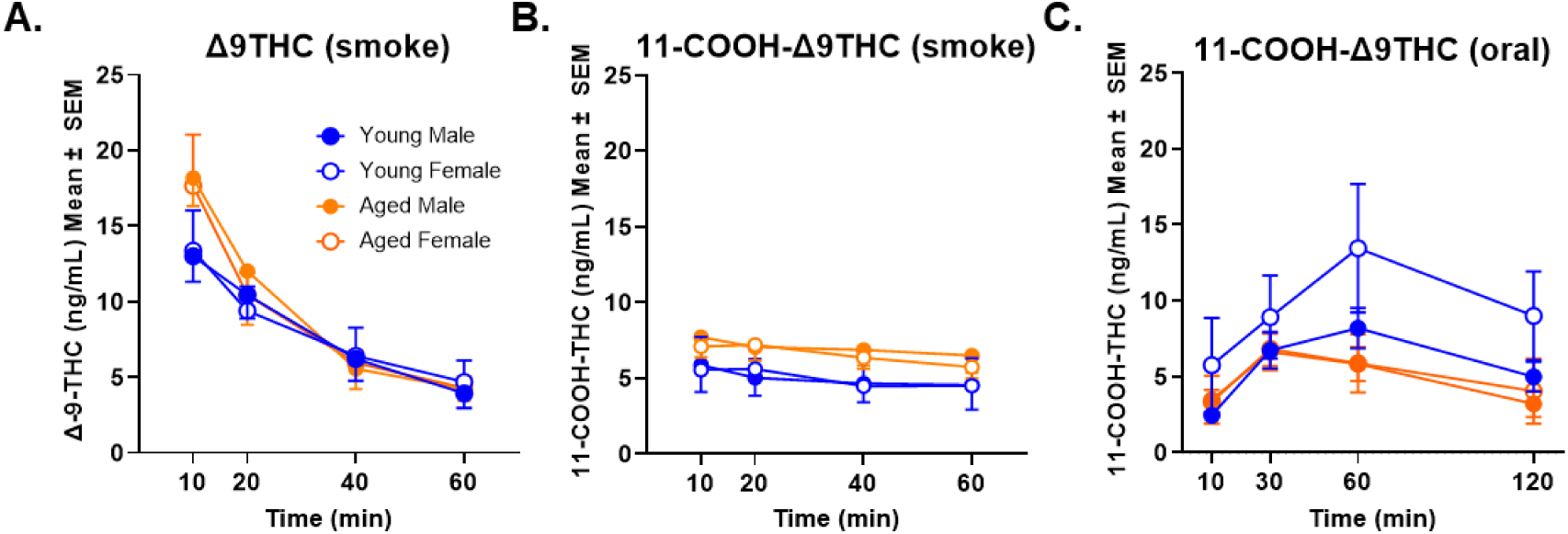
Plasma concentration-time profiles of Δ9THC and 11-COOH-Δ9THC after cannabis smoke exposure and oral administration of Δ9THC in young adult and aged rats. **A.** Peak levels of Δ9THC were evident 10 min following acute cannabis smoke exposure, but age and sex differences did not reach statistical significance (young adult: n=9 M, n=8 F; aged: n=8 M, n=12 F). **B.** Following acute cannabis smoke exposure, there were no age or sex differences in 11-COOH-Δ9THC levels. **C.** Levels of 11-COOH-Δ9THC peaked at 30-60 min following acute oral Δ9THC consumption, with levels being higher in young than aged rats, particularly at the 60-min timepoint (young adult: n=12 M, n=6 F; aged: n=10 M, n=8 F). See Table 10 for details of statistical analyses.

**Table 10.** Full statistical results for Experiment 3 (pharmacokinetics of Δ9THC following both smoke and oral administration)

| **<u>Smoke $\Delta^9$THC</u>** |  |  |
| --- | --- | --- |
| Factor | <u>F value</u> | <u>P value</u> |
| Time | **F (3,99) = 43.026** | **P<0.001** |
| Age | F (1, 33) = 0.946 | P=0.338 |
| Sex | F (1, 33) = 0.15 | P=0.903 |
| Time x Age | F (3, 99) = 2.395 | P=0.073 |
| Time x Sex | F (3, 99) = 0.262 | P=0.853 |
| Age x Sex | F (1, 33) = 0.029 | P=0.865 |
| Time x Age x Sex | F (3, 99) = 0.022 | P=0.995 |
| **<u>Smoke 11-COOH-$\Delta^9$THC</u>** |  |  |
| Factor | <u>F value</u> | <u>P value</u> |
| Time | **F (3,66) = 4.862** | **P=0.004** |
| Age | F (1, 22) = 0.823 | P=0.374 |
| Sex | F (1, 22) = 0.080 | P=0.781 |
| Time x Age | F (3, 66) = 0.170 | P=0.916 |
| Time x Sex | F (3, 66) = 0.704 | P=0.553 |
| Age x Sex | F (1, 22) = 0.980 | P=0.874 |
| Time x Age x Sex | F (3, 66) = 0.077 | P=0.972 |
| **<u>Oral 11-COOH-$\Delta^9$THC</u>** |  |  |
| Factor | <u>F value</u> | <u>P value</u> |
| Time | **F (1, 32) = 4.062** | **P=0.052** |
| Age | F (1, 32) = 3.194 | P=0.083 |
| Sex | F (1, 32) = 1.287 | P=0.265 |
| Time x Age | **F (1, 32) = 13.376** | **P<0.001** |
| Time x Sex | F (1, 32) = 1.918 | P=0.176 |
| Age x Sex | F (1, 32) = 1.133 | P=0.295 |
| Time x Age x Sex | F (1, 32) = 2.323 | P=0.137 |

**Table 11.** Numbers of samples above the LLOQ per age/sex/timepoint for each analyte.

|  |  |  |  |  |
| --- | --- | --- | --- | --- |
| **Cannabis Smoke** |  |  |  |  |
| **Δ9THC** |  |  |  |  |
| **Timepoint (Min)** | Young Male | Young Female | Aged Male | Aged Female |
| 10 | 9/9 | 8/8 | 8/8 | 12/12 |
| 20 | 9/9 | 8/8 | 7/8 | 12/12 |
| 40 | 7/9 | 7/8 | 7/8 | 12/12 |
| 60 | 5/9 | 7/8 | 6/8 | 10/12 |
| 120 | 4/9 | 2/8 | 7/8 | 3/12 |
| **11-OH-Δ9THC** |  |  |  |  |
| **Timepoint (Min)** | Young Male | Young Female | Aged Male | Aged Female |
| 10 | 3/9 | 3/8 | 1/8 | 8/12 |
| 20 | 3/9 | 3/8 | 1/8 | 7/12 |
| 40 | 2/9 | 3/8 | 0/8 | 6/12 |
| 60 | 2/9 | 3/8 | 0/8 | 7/12 |
| 120 | 1/9 | 3/8 | 0/8 | 5/12 |
| **11-COOH-Δ9THC** |  |  |  |  |
| **Timepoint (Min)** | Young Male | Young Female | Aged Male | Aged Female |
| 10 | 3/9 | 6/8 | 8/8 | 9/12 |
| 20 | 3/9 | 6/8 | 7/8 | 9/12 |
| 40 | 2/9 | 6/8 | 7/8 | 9/12 |
| 60 | 2/9 | 4/8 | 7/8 | 8/12 |
| 120 | 2/9 | 4/8 | 6/8 | 7/12 |
| **Oral Δ9THC** |  |  |  |  |
| **Δ9THC** |  |  |  |  |
| **Timepoint (Min)** | Young Male | Young Female | Aged Male | Aged Female |
| 10 | 0/19 | 2/9 | 0/15 | 1/8 |
| 30 | 4/19 | 3/9 | 2/15 | 0/8 |
| 60 | 4/19 | 1/9 | 0/15 | 1/8 |
| 120 | 1/19 | 0/9 | 0/15 | 0/8 |
| 240 | 0/19 | 0/9 | 1/15 | 0/8 |
| **11-OH-Δ9THC** |  |  |  |  |
| **Timepoint (Min)** | Young Male | Young Female | Aged Male | Aged Female |
| 10 | 1/19 | 2/9 | 0/15 | 2/8 |
| 30 | 2/19 | 4/9 | 2/15 | 3/8 |
| 60 | 1/19 | 4/9 | 0/15 | 5/8 |
| 120 | 1/19 | 4/9 | 0/15 | 2/8 |
| 240 | 1/19 | 3/9 | 1/15 | 0/8 |
| **11-COOH-Δ9THC** |  |  |  |  |
| **Timepoint (Min)** | Young Male | Young Female | Aged Male | Aged Female |
| 10 | 5/19 | 3/9 | 4/15 | 4/8 |
| 30 | 11/19 | 6/9 | 10/15 | 8/8 |
| 60 | 12/19 | 6/9 | 10/15 | 7/8 |
| 120 | 10/19 | 5/9 | 4/15 | 2/8 |
| 240 | 6/19 | 5/9 | 3/15 | 2/8 |

Following consumption of gelatin containing 1.0 mg/kg Δ9THC, plasma Δ9THC concentrations were below the LLOQ at all timepoints in all but a few rats (Table 11). Levels of 11-OH-Δ9THC were similarly low, but 11-COOH-Δ9THC was more detectable, particularly at the 30 and 60 min timepoints (Figure 7C). A three-factor ANOVA conducted on data at these two timepoints (Age x Sex x Timepoint) showed no main effect of Age (F(1,32)= 3.19, p=.08) but there was a Timepoint x Age interaction (F(1,32)= 13.38, p<.001), such that plasma 11-COOH-Δ9THC levels were higher in young adult than aged rats at the later timepoint.

## DISCUSSION

Virtually all published research in young adult subjects shows that cannabinoids impair multiple forms of cognition [23–25,27], although the doses employed are frequently outside the range of those considered to be reinforcing (e.g., [48,49]). Cannabinoid effects on cognition in older adults have been less well studied, despite this population representing the fastest-growing group of cannabis users. The growing prevalence of cannabis use among older adults in combination with their increased vulnerability to cognitive decline makes it critical to determine how cannabinoids affect cognition in this age group. Surprisingly, the few studies that have addressed this topic find no differences between older adults who use cannabis and age-matched controls on measures of attention, episodic memory, executive functions, and processing speed [32,33,50], suggesting distinct actions of cannabis on cognition across the lifespan. The present results in rats show that acute exposure to cannabis smoke enhanced working memory in aged males but impaired working memory in aged females, while having minimal effects in either sex on an episodic/spatial memory task. Chronic oral consumption of Δ9THC enhanced working memory in both sexes, while also having no effect on spatial memory. Neither drug regimen affected performance in young adults of either sex. These findings show that, at least under some conditions, cannabinoids have the potential to remediate age-related cognitive impairments.

The results of the present studies are consistent with some previous findings in aged rodents. Chronic administration of the CB_1_ receptor agonist WIN-55,212-2 via osmotic minipump enhanced water maze performance in aged rats [51]. More recently, both chronic Δ9THC (via osmotic minipump) and an acute Δ9THC injection remediated cognitive deficits across a range of tasks in aged mice [52,53]. The current findings advance this prior work in several ways. First, the routes of administration employed here mimic those used by the majority of cannabis users (including older adults [5]). As route of administration can dramatically alter Δ9THC pharmacokinetics [36], it is important to account for this variable in translating approaches and results across species. Second, unlike in prior work, subjects of both sexes were tested under the same conditions. The effects of aging on some aspects of cognition can differ in males and females (e.g., [54,55]) as can the effects of exogenous cannabinoids on cognition and behavior [56]. Thus, it is important to evaluate both sexes in determining how cannabinoids affect age-related cognitive impairments.

A notable difference between the current and previous work on cannabinoid effects in aged rodents is the absence of enhancing effects on hippocampus-mediated cognition. Whereas prior work found that Δ9THC (and the CB_1_ agonist WIN-55,232-2) improved performance on the Morris water maze in aged rodents [51–53], no such effects were observed here with chronic oral Δ9THC, or with acute cannabis smoke in the TUNL task (despite cannabis smoke being as behaviorally active in the TUNL as in the working memory task, as indexed by the reduction in number of trials completed; Figure 3). The reason for this difference is unclear (see the next paragraph for more on this point). Another notable aspect of the current findings is the fact that acute cannabis smoke impaired working memory in females, while enhancing performance in males. Δ9THC pharmacokinetics were similar in aged males and females following smoke exposure, and although there is other evidence for sex differences in cannabinoid signaling, it is unclear how they interact with advanced age [57–61]. Another possibility, however, stems from the robust sex differences in baseline performance in aged rats in Experiment 1a, such that aged females performed comparably to young adults whereas aged males were considerably impaired. Previous work in young adult rats showed that acute cannabis smoke enhances working memory only in rats (of both sexes) that performed poorly at baseline [40], suggesting that baseline performance rather than sex accounts for the sex difference in acute cannabis smoke effects in aged rats in this experiment. This is supported by the results of Experiment 2, in which aged control rats of both sexes were impaired relative to young, and both benefitted from chronic Δ9THC.

In considering the translational implications of these findings, it is important to consider cannabis use patterns amongst older adults. Many older adults report using cannabis regularly to treat chronic medical conditions, with edibles (49%) and smoking (29%) being the most common routes of intake [43,62]. In addition, the doses of Δ9THC ingested through both smoke and oral routes in this study were relatively low, and although behaviorally active ([41,42] and current data), they likely produced limited psychoactive effects (e.g., rats did not show a taste aversion to oral Δ9THC). These low doses could account for the absence of effects on hippocampus-dependent cognition compared to previous work [51–53,63], although the different drugs, routes of administration, and species employed render such comparisons challenging. Importantly, however, the levels of Δ9THC exposure in the present study are well below those shown to produce cognitive impairments in young rats, including in both working memory and water maze tasks [40,48,64,65] (see also [66] for related findings). Finally, note that exposure to smoke alone (aside from its Δ9THC content) was unlikely to have contributed to its cognitive effects, as exposure to smoke from placebo cigarettes (from which cannabinoids are removed) does not mimic effects of cannabis smoke on cognition [40,67].

Δ9THC exposure in aged mice causes a host of changes in the hippocampus, including increases in dendritic spine density and synaptic protein expression that are linked to enhanced hippocampus-dependent cognition [35,68]. In contrast, the mechanisms by which cannabis/ Δ9THC might enhance working memory in aging are less clear. Performance on delayed response tasks is thought to depend upon sustained PFC pyramidal neuron activity, which is regulated by input from GABAergic interneurons [69]. Age-related working memory impairments are linked to alterations in PFC pyramidal neuron excitability, due in part to increased GABAergic tone [70–72]. CB_1_ receptors are highly expressed on PFC GABAergic interneurons [60,73–75], and acute activation of these receptors could suppress GABA release, thus disinhibiting pyramidal neuron activity [76]. Such pyramidal neuron disinhibition could provide a mechanism by which cannabis acutely improves PFC-dependent working memory. In contrast to PFC, age-related impairments in hippocampus-mediated cognition in both rats and humans are due in part to hyperexcitability of pyramidal neurons in the CA3 subregion [77,78]. Given that CB_1_Rs are also highly expressed by a subset of hippocampal GABAergic interneurons [79], Δ9THC-induced suppression of GABAergic activity might exacerbate hippocampal excitability, and provide more limited cognitive benefits in aging.

It is less likely that acute Δ9THC actions on CB_1_Rs account for the enhancing effects on working memory in Experiment 2. Although circulating levels of the secondary Δ9THC metabolite 11-COOH-Δ9THC were higher in young adult than aged rats (and particularly in females, consistent with [46,80,81]), these levels would have been extremely low at the time of cognitive testing 20 h after oral administration (Figure 7 and Table 11). An alternative mechanism concerns Δ9THC actions on inflammatory signaling. Cognitive impairments accompanying aging are associated with dysregulated inflammatory signaling and increased circulating levels of pro-inflammatory cytokines [82–90]. Within the brain, pro-inflammatory factors can act on both microglia and neurons via specific receptors, and can ultimately modulate neuronal excitability [86,91–96]. Δ9THC has anti-inflammatory effects both *in vitro* and *in vivo*, including attenuation of microglial activation and suppression of pro-inflammatory cytokine expression [97–99], suggesting that chronic Δ9THC could attenuate age-related increases in inflammatory signaling and ultimately remediate cognitive deficits [51,63].

The results of this study demonstrate that cannabis does not impair, and can provide benefit to, cognition in older subjects, but that these benefits vary as a function of sex, duration of exposure, and/or the form of cognition being measured. It is also important to highlight that the route of administration between acute and chronic studies differed, and that their distinct pharmacokinetic profiles could impact the effects of cannabinoids on cognition as a function of sex or age, rendering direct comparisons between the two experiments challenging. In addition, we did not assess potential age differences in the pharmacokinetics of Δ9THC in brain. A further factor to consider is that the experiments evaluated effects of Δ9THC alone, and did not investigate contributions of other cannabis constituents such as cannabidiol, which can interact with the effects of Δ9THC on cognition and other aspects of behavior [34,100,101]. Future work will assess effects of cannabidiol on cognition in the context of aging. Moving forward, it will be important to begin identifying mechanisms driving the pro-cognitive effects of cannabis in aging.

## MATERIALS AND METHODS

### Subjects

Fischer 344 x Brown Norway F1 hybrid (FBN) rats obtained from the National Institute on Aging colony maintained by Charles River were used across all experiments. Rats were 6-9 months (young adult) and 24-28 months (aged) at the start of experiments. Rats were single housed in polycarbonate cages (38 cm × 28 cm × 20 cm) containing Sanichip bedding (P.J. Murphy Forest Products) and a wire top, and maintained on a reversed 12-h light/dark schedule (lights off at 0800). Upon arrival, rats were given one week to acclimate prior to initiating any procedures. All experiments took place during the dark phase, 5-7 days per week, at approximately the same time each day. For the food-motivated cognitive tasks, rats were placed on a food restriction plan. Food consisted of standard rat diet and was limited to attain a maintenance weight of 80-85% of their free-feeding weight. Throughout the study, rats were weighed daily, and body condition score was assessed and recorded weekly to maintain a range of 2.5-3.5. The body condition score was assigned based on the presence of palpable fat deposits over the lumbar vertebrae and pelvic bones [102,103]. Water was provided in the home cage *ad libitum*. All animal studies were performed in accordance with the University of Florida Institutional Animal Care and Use Committee and National Institutes of Health guidelines.

### Experiment 1: Effects of cannabis smoke on cognition in young adult and aged rats

Experiment 1 was designed to determine how acute exposure to cannabis smoke affects cognitive performance in young adult and aged rats of both sexes. Rats were trained on a delayed response working memory task and trial-unique non-match to location (TUNL) task until stable performance emerged, followed by test sessions in which the effects of acute smoke exposure were evaluated using a within-subjects design. Some rats were trained and tested in both behavioral tasks, with the order of tasks counterbalanced across age and sex.

#### Smoke exposure apparatus and procedures

Smoke exposure was conducted in an automated cigarette smoking machine (model TE-10, Teague Enterprises, Davis, CA) as described previously [41,46]. During smoke exposure sessions, rats were pair housed in clean polycarbonate cages (38 × 28 × 20 cm) containing Sanichip bedding and a wire top. Four cages were placed into the exposure chamber of the machine for each smoke session. Cannabis cigarettes were placed in the ignition chamber, where they were lit and puffed (35 cm^3^ puff volume, 1 puff per min, 2 s per puff). Smoke from the ignition chamber was distributed evenly throughout the exposure chamber and exhausted to the building exterior. Each cigarette required approximately 10 puffs to burn completely (10 min total), after which it was replaced with a fresh cigarette and the process repeated. At the completion of smoke exposure, there was a 15 min purge period to evacuate smoke from the exposure chamber, after which rats were removed and immediately began cognitive testing.

#### Drug

Cannabis cigarettes (approximately 700 mg each) were obtained from the National Institute on Drug Abuse Drug Supply Program, and contained approximately 5.6% Δ9-tetrahydrocannabinol (Δ9THC), 0% cannabidiol (CBD), and 0.4% cannabinol (CBN) as per the accompanying analytical data. Cigarettes were stored at −20° C until 24 hours prior to use, when they were transferred to a humidifier chamber. Rats were exposed to smoke from 0, 3, and 5 consecutively-burned cigarettes using a randomized, within-subjects design such that each rat was tested in separate sessions under each exposure condition. At least 48 h elapsed between consecutive exposure sessions.

#### Behavioral Testing

##### Apparatus

Behavioral testing was conducted in eight identical Bussey-Saksida rat touchscreen operant chambers (Lafayette Instruments, Lafayette, IN) equipped with a touch-sensitive monitor (38.1 cm, screen resolution 1024 horizontal pixels x 768 vertical pixels), a fan (providing ventilation and white noise), a house light, and a food trough (with a light to illuminate the trough and an infrared beam to detect head entries). A single 45 mg food pellet (AIN-76A, Test Diet, Richmond, IN) delivered into the food trough was used to reinforce correct responses. The operant chambers consisted of black Perspex walls that formed an isosceles trapezoid-shaped floor, with a transparent lid and a grated metal floor. The chambers were housed within sound-attenuating cabinets to minimize outside noise. During training, a black plastic mask was used to divide the touchscreen into two rows of 7 squares (each 2 cm x 2 cm), but only the bottom row of 7 squares was utilized for behavioral testing (see Figures 1A and 2A). This mask created distinct response windows to guide rats in interacting with the screen. Stimulus presentation and data collection were controlled by computers running ABET II Touch software (Campden Instruments Ltd, Lafayette, IN) and Whisker Server [104]. Prior to the start of both behavioral tasks, rats went through a series of shaping procedures designed to train them to perform the various task elements (see [105] for detailed descriptions of shaping). The shaping protocol consisted of five 1-h stages. In Stage 1 (“magazine training”) a single food pellet was dispensed following a nosepoke into the food trough. In Stage 2 (“full screen any touch”) the entire touchscreen was illuminated, and rats received a single pellet for touching anywhere on the screen. In stage 3 (“progressive any touch”), the entire screen was illuminated for the first 20 trials, and then only the top three-quarters of the screen were illuminated for the next 20 trials, and finally the top half of the screen was illuminated for the remainder of the session. This stage of shaping helped gradually guide rats’ attention to the upper portion of the screen where relevant stimuli would be presented during behavioral testing. In Stage 4 (“any touch”) a mask was applied over the screen such that rats could only interact with the small square cutout panels in the mask. Multiple panels were illuminated on the screen on each trial, and rats received a single pellet for touching any of them. In Stage 5 (“must touch”) only a single panel was illuminated on the screen, and rats received a single pellet only after touching that panel. To prevent formation of positional biases, a different panel was illuminated on each trial. In Stage 6 (“must initiate”), rats were required to nosepoke into the (illuminated) food trough to trigger illumination of one of the panels (once the panel was illuminated, the contingencies were identical to those in Stage 5). Finally, in Stage 7, (“punish incorrect”), failure to touch the illuminated panel resulted in the houselights being turned on, and a 10-s timeout period began before rats were allowed to initiate the next trial.

##### Working memory task

The delayed response working memory task consisted of daily 60-min sessions composed of multiple trials, which were initiated with a nosepoke into the illuminated food trough. Each trial was divided into a sample phase, a delay phase, and a choice phase. During the sample phase, one panel on the touchscreen display (either the far left or the far right, randomized across trials) was illuminated (Figure 1A). A touch upon the illuminated panel caused the screen to go blank and initiated the delay phase, which was of a duration of 0, 6, 12, or 24 s (randomly selected from within blocks of four trials that contained one trial at each delay). During the delay phase, the food trough, which was located on the wall opposite from the touchscreen, was illuminated, and the rat was required to nosepoke into the trough. After the delay elapsed, the choice phase began, during which rats were presented with two illuminated panels (the sample and the symmetrically-placed panel on the opposite side of the screen). Rats were required to select the panel that was not previously illuminated during the sample phase (i.e., employ a non-match rule) in order to receive a food pellet. Selection of the “incorrect” panel caused the screen to go blank and the houselight to be illuminated, and no food was delivered. Upon completion of the choice phase, a 5 s intertrial interval elapsed before the food trough was illuminated and the next trial could be initiated.

##### Trial-Unique Non-matching to Location Task

The TUNL task consisted of daily 60-min sessions composed of multiple trials, which were initiated with a nosepoke into the illuminated food trough. Each trial was subdivided into a sample phase and a choice phase. During the sample phase, one panel on the touchscreen display (randomly selected from among the bottom 7 positions) was illuminated (Figure 2A). A touch upon the illuminated panel caused the screen to go blank and the light in the food trough was illuminated. A nosepoke into the food trough caused two panels on the touchscreen to be illuminated (both the original location and a second location separated by 1, 3, or 5 blank locations, randomly selected from within blocks of three trials that contained one trial at each separation). Rats were required to touch the panel that was not illuminated during the sample phase (i.e., employ a non-match rule) in order to receive a single food pellet. A touch on the “incorrect” panel caused the screen to go blank and the house light to be illuminated, and no food was delivered. Upon completion of the choice phase, a 5-s intertrial interval elapsed before the food trough was illuminated and the next trial could be initiated.

#### Data analyses

Raw behavioral data files were exported from ABET software into Microsoft Excel and analyzed further using SPSS 28.0. The primary measure of performance in both the delayed response and TUNL tasks was accuracy (percentage of correct trials at each delay or stimulus separation, averaged over the 60-min session). Prior to smoke exposure, rats were trained on their respective task until reaching a stable baseline of performance. Stability was defined as the absence of a main effect or interaction involving session in a two-factor, repeated measures analysis of variance (ANOVA; Delay or Separation X Session) conducted on data from three consecutive sessions. For both the delayed response and TUNL tasks, exclusion criteria were applied prior to data analysis to remove rats with poor task engagement under control conditions or insufficient data under any condition. For the delayed response task, the exclusion criteria were less than 80% accuracy at the 0-s delay in the 0-cigarette condition (indicating poor learning of task rules) or fewer than 35 trials in a session under any cigarette condition. A total of n=9 Aged and n=1 Young rats were excluded for the Delayed response task analysis. For the TUNL task, the exclusion criteria were fewer than 30 trials per session under any cigarette condition. A total of n=7 Aged and n=1 Young rats were excluded for TUNL task analysis. To assess the effects of cannabis smoke on each task, data were initially analyzed using four-factor, repeated measures ANOVAs (Age x Sex x Cigarette condition x Delay or Separation), with Cigarette condition and Delay or Separation as within-subjects variables. The initial ANOVAs were followed up by two-factor ANOVAs within each age and sex. The numbers of completed trials per session were analyzed via three-factor repeated measures ANOVA (Age x Sex x Cigarette condition), with Cigarette condition as the within-subjects factor. In all cases, p less than or equal to .05 were considered statistically significant.

### Experiment 2: Effects of chronic oral Δ9THC on cognition in young and aged rats

Experiment 2 was designed to determine how chronic oral consumption of Δ9THC affects cognitive performance in young adult and aged rats. Rats were trained on a delayed response working memory task until stable performance emerged, followed by division into two groups matched for performance, and further testing on the task under Δ9THC or control conditions for 3 weeks. This was followed by testing in the Morris water maze under the same conditions for a further two weeks.

#### Drug administration procedures

Δ9THC was obtained from the NIDA Drug Supply Program in ethanol stock solution. The stock solution was diluted using 95% ethanol to a working concentration of 10 mg/ml, and was stored at −20° C under nitrogen gas. Δ9THC was administered in gelatin form, which was presented to the rats in their home cages. To make the gelatin, 3.85 g of Knox Gelatin was dissolved in 100 ml of deionized water and raised to a temperature of 90° C, followed by addition of 5 g of Polycal sugar. Gelatin was cooled at 4° C before use. To provide the gelatin to the rats, 10 g of gelatin was placed in plastic feeding containers (Nombrero, Animal Specialties and Provisions, Quaker Town, PA), which were secured to the cage lids. Gelatin exposure began with 5 sessions (one session/day) designed to acclimate rats to the gelatin and overcome neophobia. Just prior to placing the gelatin in the home cage, a small amount of 95% ethanol (100 µl/kg rat weight) was added to the gelatin in the feeder. After 1 h of exposure, the remaining weight of the gelatin was recorded. At this point, rats of each age and sex were divided into two groups (Δ9THC and control), matched for baseline working memory accuracy. The control group continued to receive gelatin with ethanol, while the Δ9THC group received gelatin with Δ9THC added (100 µl/kg of a 10.0 mg/ml solution, for a dose of 1.0 mg/kg). Behavioral testing was conducted in the mornings, and gelatin administration was conducted in the afternoons, in order to eliminate potential acute effects of Δ9THC consumption. Both testing and gelatin administration were conducted Monday-Friday.

#### Behavioral Testing

##### Apparatus

###### Operant chambers

The delayed response working memory task was conducted in 8 identical operant test chambers (30.5 cm x 25.4 cm x 30.5 cm) with Plexiglas side walls and stainless steel front and rear walls, which were housed in sound-attenuating cabinets (Coulbourn Instruments, Holliston, MA). The floor of each chamber was composed of stainless steel rods, and a food pellet delivery trough was located on the front wall. The trough was 2 cm above the chamber floor and contained a photobeam to detect nosepoke entries as well as a 1.12 W lamp. Two retractable levers placed 11 cm from the floor were located on either side of the trough. A 1.12 W house light was mounted on the rear wall of the sound-attenuating cabinet. The operant chambers interfaced with a computer running Graphic State 4.0 software (Coulbourn Instruments) to allow for experiment control and data collection.

###### Water maze

The maze consisted of a circular tank (diameter 183 cm, wall height 58 cm) painted white and filled with water (27° C) made opaque with the addition of non-toxic white tempera paint. A retractable escape platform (12 cm diameter, HVS Image, UK) was submerged 2 cm below the water’s surface near the center of one quadrant of the maze (spatial learning task) or protruded 2 cm above the water’s surface and was moved to a different maze quadrant on each trial (cued learning task). The maze was surrounded by black curtains, to which were affixed large white geometric designs that provided extramaze cues. Swim data were recorded via an overhead video camera, and analyzed using a computer-based video tracking system (Water 2100, HVS Image, UK).

##### Working memory task

This version of the delayed response working memory task began with shaping procedures designed to train rats to perform the various task components. Shaping Stage 1 was magazine training. In this 64-min protocol, rats were required to reach a total of 100 nosepokes into the food trough, into which food pellets were delivered on a variable interval schedule of 100 +/− 40s. After reaching criterion on Stage 1, rats moved to Shaping Stage 2 (single lever training), in which they were trained to press one of the levers (either left or right, counterbalanced across groups) to earn a single food pellet. Pressing the lever at least 50 times in the span of a 30-min session progressed them to a shaping session with the other lever under the same criterion. Once rats reached criterion on both levers, they proceeded to Shaping Stage 3, which consisted of multiple trials on which a single lever (left or right, randomized within pairs of trials) was extended into the chamber, and a press earned a single food pellet. Criterion in this stage was at least 30 presses on each lever in the 60-min session. Completion of Stage 3 moved the rats on to Shaping Stage 4, in which a nosepoke into the food trough initiated extension of one of the two levers. A press on this lever earned a single food pellet. Rats were trained in this stage for 1 h/day until they obtained 80 lever presses in a session for a minimum of four sessions. Rats then proceeded to the delayed response working memory task.

The delayed response working memory task was modified from [37] and consisted of multiple trials within each 40-min session. Each trial began with a single lever (the “sample” lever) being extended into the operant chamber (Figure 4A). The left/right position of the sample lever was randomized within pairs of trials, and the lever was retracted once pressed, which initiated the “delay” phase. During the delay, rats were required to nosepoke into the food trough, in order to minimize the use of mediating strategies such as remaining in front of the sample lever. The first nosepoke after the delay period expired initiated the “choice” phase, in which both levers were extended. A press on the same lever extended in the sample phase (the “correct” lever) resulted in delivery of a food pellet (i.e., rats had to employ a “match-to-sample” rule). A press on the opposite (“incorrect”) lever caused the levers to retract, the houselight to extinguish, and no food delivery. Both correct and incorrect responses were followed by a 5-s intertrial interval, after which the next trial began. The house light was illuminated throughout the session except during the intertrial intervals following incorrect responses.

The durations of the delay phase between the sample and choice phases of the trials were increased successively over the course of training. Initially, the delay duration was set at 0 s. Correction trials were used during this stage, such that if rats pressed the incorrect lever they were presented with the same sample lever on the next trial. Once rats achieved greater than 80% accuracy over 2 consecutive sessions on a set of delays, the delay durations were increased (delay set 1: 0, 1, 2, 3, 4, 5, 6 s; delay set 2: 0, 2, 4, 8, 12, 16 s; delay set 3: 0, 2, 4, 8, 12, 18, 24 s – note that correction trials were discontinued for the three sets with non-zero delays). Within each set of delays, the order of the delays was randomized within blocks of 7 trials. Delay set 3 was used for all behavioral testing reported in the manuscript [13,106].

##### Morris water maze task

The water maze protocol was based on [18,44,45] To assess spatial learning and memory, rats received three swim trials/day over eight consecutive days, with a 60-s intertrial interval. On each training trial, rats were placed in the water facing the wall of the maze and allowed to swim until finding the escape platform (which remained in the same quadrant for all 8 days) or until 90 s elapsed, at which point they were guided to the platform by the experimenter. Rats were allowed to remain on the platform for 30 s before removal from the maze and the start of the intertrial interval. The starting position for each trial varied pseudorandomly among four equally spaced positions around the perimeter of the maze (north, south, east, and west). Every sixth trial was a probe trial on which the escape platform was removed, and rats swam for the full 90-s trial.

To assess rats’ sensorimotor abilities and motivation to escape the water independent of spatial learning, rats received one session with six trials of cued water maze training following the last day of hidden platform (spatial) training. In this session, rats were trained to escape to a visible black platform that protruded 2 cm above the water’s surface and which was moved to a different maze quadrant on each trial. On each trial, rats were given 90 s to reach the platform and were allowed to remain there briefly before a 30-s intertrial interval.

#### Data analyses

In the working memory task, raw datafiles were exported from Graphic State 4.0 into Microsoft Excel, and subsequently analyzed using SPSS 28.0. The primary measure of performance in this task was accuracy (percentage of correct trials at each delay, averaged over the 40-min session). Prior to gelatin exposure, rats were trained until reaching stable performance, defined as the absence of a main effect or interaction involving session in a two-factor, repeated measures ANOVA (Delay X Session) conducted on data from three consecutive sessions. To assess the effects of chronic THC consumption, data from week 3 of testing were initially analyzed using a four-factor, repeated measures ANOVA (Age x Sex x Drug condition x Delay), with Delay as a within-subjects variable. The initial ANOVA was followed up by three- or two-factor ANOVAs when warranted. The number of completed trials was analyzed via three-factor ANOVA, with Age, Sex, and Drug condition as between-subjects variables.

In the water maze, data were exported from HVS Image software to Microsoft Excel and analyzed using SPSS 28.0. The primary performance measure in the spatial learning task was search error, calculated by sampling the rat’s distance from the platform 10 times/s and averaging this distance into 1-s bins. On training trials, a cumulative search error measure (sum of the 1-s distances from the platform across the entire duration of the trial) was used, whereas on probe trials, a mean search error measure (average of distances from the platform across the entire 30-s trial) was used. On training trials in the cued task, the swim pathlength to reach the platform, averaged across trials 2 through 6, was used as the measure of performance. Rats with cued task pathlengths > 2 standard deviations from the mean of the young group were excluded from water maze analyses. Two young and one aged rat met this exclusion criterion and were excluded from subsequent analyses. For all three measures, data were analyzed using a four-factor, repeated measures ANOVA, with Spatial training trial block, Probe trial, or Cued training trial as the within-subjects variables, and Age, Sex, and Drug condition as between-subjects variables, with follow-up three- or two-factor ANOVAs as appropriate.

### Experiment 3: Pharmacokinetics of Δ9THC and its metabolites following cannabis smoke exposure and oral Δ9THC consumption

#### Cannabis smoke exposure

Rats were exposed to smoke from burning 5 cannabis cigarettes as described in Experiment 1. This was followed by a 15-min purge period to evacuate smoke from the exposure chamber, after which rats were removed from the smoke exposure apparatus (this removal point was considered “time 0”). Blood was collected at 10, 20, 40, 60, and 120 min timepoints and evaluated for concentrations of Δ9THC and two major metabolites (11-OH-Δ9THC and 11-COOH-Δ9THC) via ultraperformance liquid chromatography tandem mass spectrometry (UPLC-MS/MS) as in [4,13].

#### Oral consumption of Δ9THC

Rats were allowed to consume gelatin containing 1.0 mg/kg Δ9THC as described in Experiment 2, followed by blood collection at 10, 30, 60, 120, and 240 min timepoints and evaluation of Δ9THC, 11OH-Δ9THC, and 11COOH-Δ9THC via UPLC-MS/MS.

##### Blood collection

Serial blood samples of 200 µl of whole blood were collected via tail nicks into heparin-coated microtubes. Plasma (50 µl) was separated by centrifuging at 6500 rpm for 15 min, transferred to a 0.6 ml polypropylene microcentrifuge tube, and frozen at −80°C until analysis.

##### Analysis of Plasma Δ9THC and metabolites

###### Materials and reagents

Commercially-available standards (purity >98%) for Δ9THC and its 11-COOH-Δ9THC and 11-OH-Δ9THC metabolites were obtained from Cerilliant (Round Rock, TX, USA). Liquid chromatography–mass spectrometry (LC-MS)-grade water, acetonitrile, isopropanol, methanol, and formic acid were obtained from Fisher Scientific (Fair Lawn, NJ). Blank rat plasma was obtained from Fisher Scientific.

###### Instrumentation and analytical conditions

The UPLC-MS/MS bioanalytical method used to analyze samples was developed based on a previously validated method for use of detection of cannabinoids in low Δ9THC varieties of cannabis [13]. A Waters Acquity Class-I chromatograph coupled with a Xevo TQ-S Micro triple quadrupole mass spectrometer was used for separation and detection of cannabinoids and their metabolites. The system was controlled by MassLynx 4.2 with data processed using TargetLynx XS (Waters, Milford, MA). Chromatographic separation was achieved on a Waters Acquity UPLC BEH C18 column (1.7 µm, 2.1 × 100 mm) with a VanGuard precolumn of the same chemistry. The mobile phase consisted of water acidified with 0.1% formic acid and 50:50 (v:v) methanol and acetonitrile at a flow rate of 0.35 mL/min. The gradient elution started at 18% A which was linearly decreased to 0% over 4.5 min before a steep return to the initial conditions for re-equilibration of the column. Column and autosampler temperatures were maintained at 40° C and 10°C, respectively. The eluent was analyzed in positive electrospray ionization mode using multiple reaction monitoring. The capillary voltage was 0.5 kV, the source temperature was 150° C, the desolvation temperature was 450° C, the desolvation gas flow was 800 L/h, and the cone gas flow was 60 L/h.

###### Preparation of calibration and quality control standards

Two mix stock solutions (10000 ng/ml and 1000 ng/ml for each analyte) were prepared by combining the appropriate volume of 100 µg/ml primary stocks. These mix stocks were used to make eight working stock solutions (25, 50, 100, 250, 500, 1000, 1500, 2500 ng/mL). Each calibration standard was prepared by spiking 20 µl of blank plasma with 2 µl of the associated working stock to yield plasma concentrations in the range of 2.5-250 ng/ml. The quality control samples were prepared from a different primary stock using the same method at the four following concentrations: 2.5 ng/ml (lower limit of quantification (LLOQ)), 6 ng/ml (low quality control (LQC)), 80 ng/ml (medium quality control (MQC)), and 180 ng/ml (high quality control (HQC)).

###### Sample preparation

Samples were thawed to room temperature immediately prior to analysis. Each sample was mixed by vortex and 20 µl was subjected to a simple and fast protein precipitation method for the removal of endogenous substances and extraction of compounds of interest. Plasma samples, calibration standards, and quality control samples were quenched with 100 µl of methanol containing 0.05% formic acid and 10 ng/mL internal standard. The samples were vortex mixed for 5 min at 650 rpm before being transferred to a 96-well Millipore (Burlington, MA) multiscreen Solvinert 0.45 µm filter plate. The samples were filtered by centrifugation at 805×g for 2 min at 4°C. The filtrate was then subjected to analysis.

##### Statistical Analyses

Plasma concentrations of Δ9THC, 11-OH-Δ9THC, and 11-COOH-Δ9THC across timepoints were expressed in ng/ml. For some of the analytes and/or sampling timepoints, however, many values were below the lower limit of quantification (LLOQ, 2.5 ng/ml) (see Table 11), rendering complete analyses challenging. To analyze these data, first, rats that had no values above the LLOQ at any timepoint were removed from the analyses altogether (the large numbers of rats removed for this reason precluded analysis of 11-OH-Δ9THC for the smoke condition, and both Δ9THC and 11-OH-Δ9THC for the oral Δ9THC condition). Next, for the remaining rats, only analytes and timepoints for which at least 90% of the values were above the LLOQ were subjected to statistical analyses. For these analyses, values below the LLOQ were replaced with half of this value (1.25 ng/ml) and analyzed via three-factor, repeated measures ANOVA, with Age and Sex as between-subjects variables and Timepoint as a within-subjects variable.

## ACKNOWLEDGEMENTS

We thank the Drug Supply Program at the National Institute on Drug Abuse for providing cannabis cigarettes and THC, and Bonnie McLaurin, Vicky Kelley, Brandon Hellbusch, Bailey McCracken, and Matthew Bruner for their technical assistance with the project. Behavioral schematics in figures were created using Biorender. This work was supported by NIH R01AG072714 (BS, JLB), NIH T32AG061832 (SZ), the Florida Department of Health Ed and Ethel Moore Alzheimer’s Disease Research Program grant 21A11 (BS, JLB), and the McKnight Brain Research Foundation (JLB).

